# Developmental innovations promote species diversification in mushroom-forming fungi

**DOI:** 10.1101/2021.03.10.434564

**Authors:** Torda Varga, Csenge Földi, Viktória Bense, László G. Nagy

## Abstract

Fungi evolved complex fruiting body (‘mushroom’) morphologies as adaptations to efficient spore dispersal in terrestrial habitats. Mushroom-forming fungi (Agaricomycetes) display a graded series of developmental innovations related to fruiting body morphology, however, how these evolved is largely unknown, leaving the functional biology and evolutionary principles of complex multicellularity in the third largest multicellular kingdom poorly known. Here, we show that developmental innovations of mushroom-forming fungi that enclose the spore-producing surface (hymenophore) in a protected environment display significant asymmetry in their evolution and are associated with increased diversification rates. ‘Enclosed’ development and related tissues (partial and universal veils) evolved convergently and became a widespread developmental type in clades in which it emerged. This probably mirrors increased fitness for protected fruiting body initials in terrestrial habitats, by better coping with environmental factors such as desiccation or predators, among others. We observed similar patterns in the evolution of complex hymenophore architectures, such as gills, pores or teeth, which optimize biomass-to-propagule number ratios and were found to spur diversification in mushrooms. Taken together, our results highlight new morphological traits associated with the adaptive radiation of mushroom-forming fungi and present formal phylogenetic testing of hypotheses on the reproductive ecology of a poorly known but hyperdiverse clade.

## Introduction

Increasing reproductive efficiency is of prime importance to all organisms and has prompted the evolution of sophisticated mechanisms for protecting offspring. Diverse solutions evolved for protecting developing youth across the tree of life; all these share nursing and protective mechanisms that optimize the nutritional investment of the individual per propagulum. Examples include placentation (Roberts et al., 2016), viviparity and matrotrophy (Blackburn, 1999) in animals or the seed in embryophytes (Goldberg et al., 1994). Many such traits are considered key innovations that have spurred lineage diversification (e.g., viviparity in fishes, Helmstetter et al., 2016), have arisen convergently (e.g., viviparity occurred ∼150 times in vertebrates, Blackburn, 2015) or underline the evolutionary success of diverse clades (e.g., seed plants, Westoby & Rice, 1982, but see Vamosi et al., 2018).

Fungi reproduce by sexual or asexual spores, which are born on specialized spore-producing cells. In mushroom-forming fungi (Agaricomycetes), these cells compact into a spore producing surface, the hymenophore, in which meiosis, spore production and dispersal takes place. The hymenophore is exposed to environmental impacts (e.g., desiccation, precipitation, UV radiation), fungivorous animals, and parasites and many strategies evolved to protect the hymenophore from these (e.g., Braga et al., 2015). One such solution is the development of complex fruiting bodies, which provide support, physical barrier and chemical defense against external factors (Künzler, 2018) as well as facilitates spore dispersal (Dressaire et al., 2016). Physical protection comes in many forms, including hyphal sheaths that cover either the entire fruiting body initial (universal veil) or parts of it (partial veil), or producing spores inside the fruiting body (in gasteroid and secotioid fungi). All these strategies enclose the hymenophore into a protected environment, at least during early developmental stages, and we hereafter refer to it as enclosed development.

Several key principles of the evolution of fruiting bodies have been uncovered recently. Phylogenetic comparative analyses confidently suggest that ancestral morphologies were crust-like and that these repeatedly gave rise to a series of more complex forms. The most derived ones are called pileate-stipitate morphologies (mushrooms with cap and stalk), which evolved several times convergently and probably represent stable attractors in the morphospace (Hibbett, 2004, Varga et al. 2019). Further, the emergence of complex morphologies correlate with higher diversification rates and may be a major driver of lineage diversification in mushroom-forming fungi (Agaricomycetes) (Sánchez-García et al., 2020; Varga et al., 2019). However, beyond the broadest morphological types, we know little about what drives the evolution of fruiting body morphologies and how novel fruiting body traits impact speciation and extinction patterns. For example, it is not known what aspect of the pileate-stipitate morphology – protection of the hymenophore, increased efficiency of spore dispersal or yet other attributes – may have been the key innovation for mushroom-forming fungi. Further, there are several phylogenetically co-distributed morphological innovations, such as structured hymenophore surfaces, which could additively or in other ways influence diversification rates.

Here, we investigate the evolution of enclosed development among mushroom-forming fungi using comparative phylogenetic analyses and a previously published phylogeny of 5,284 species (Varga et al., 2019). We demonstrate that enclosed development evolved repeatedly in the Agaricomycetes and correlates to increased diversification rate of species. We further show that other, phylogenetically co-distributed traits (complex hymenophores, the presence of a cap) also impact diversification rates, but their effects are independent from those of enclosed development. Our results reveal novel factors in the adaptation of mushroom-forming fungi to terrestrial habitats.

## Material and Methods

### Phylogenetic data

All macro-evolutionary analyses were performed on 245 Maximum likelihood phylograms and ten chronograms inferred in our previous work (Varga et al., 2019). These trees were inferred from three loci (28S subunit ribosomal RNA, ef1-alpha and RPB2) and a phylogenomic backbone tree of 104 species, and represents a robust evolutionary framework with ca. one-fifth of all described species in Agaricomycetes sampled. Species from the classes Dacrymycetes and Tremellomycetes were used as an outgroup. Time calibration of trees was performed in a two-step Bayesian analysis on ten randomly sampled phylogenies.

### Character coding

#### Developmental types

The character state assignment was based on whether the developing hymenophore is open to the environment or insulated from it at some point during development (figure 1, table S1). In our default coding regime (referred to as 3ST), we divided the diversity of developmental types into three character states, open (state 0), semi-enclosed (state 1) and enclosed development (state 2). Open development was defined as the hymenophore being exposed to the environment from the earliest primordial stages and corresponds to gymnocarpy *sensu* Reijnders (1948) or exocarpy without any metablemas *sensu* Clémencon (2012). In semi-enclosed development, the hymenophore is covered by a veil (usually faint) only in the earliest primordial stages or the cap margin is attached to the stem but detaches before the start of the cap expansion (hypovelangiocarpy and pilangiocarpy *sensu* Reijnders (1948)). In enclosed development, the hymenophore is closed at least until the young fruiting body stage (angiocarpy *sensu* Reijnders (1948) or endocarpy and nodulocarpy *sensu* Clémencon (2012)). We coded gasteroid/secotioid species as enclosed. For historical reasons, this morphology is often treated as a separate state, however, from the perspective of this character, they represent a special case of enclosed development.

**Figure 1.**
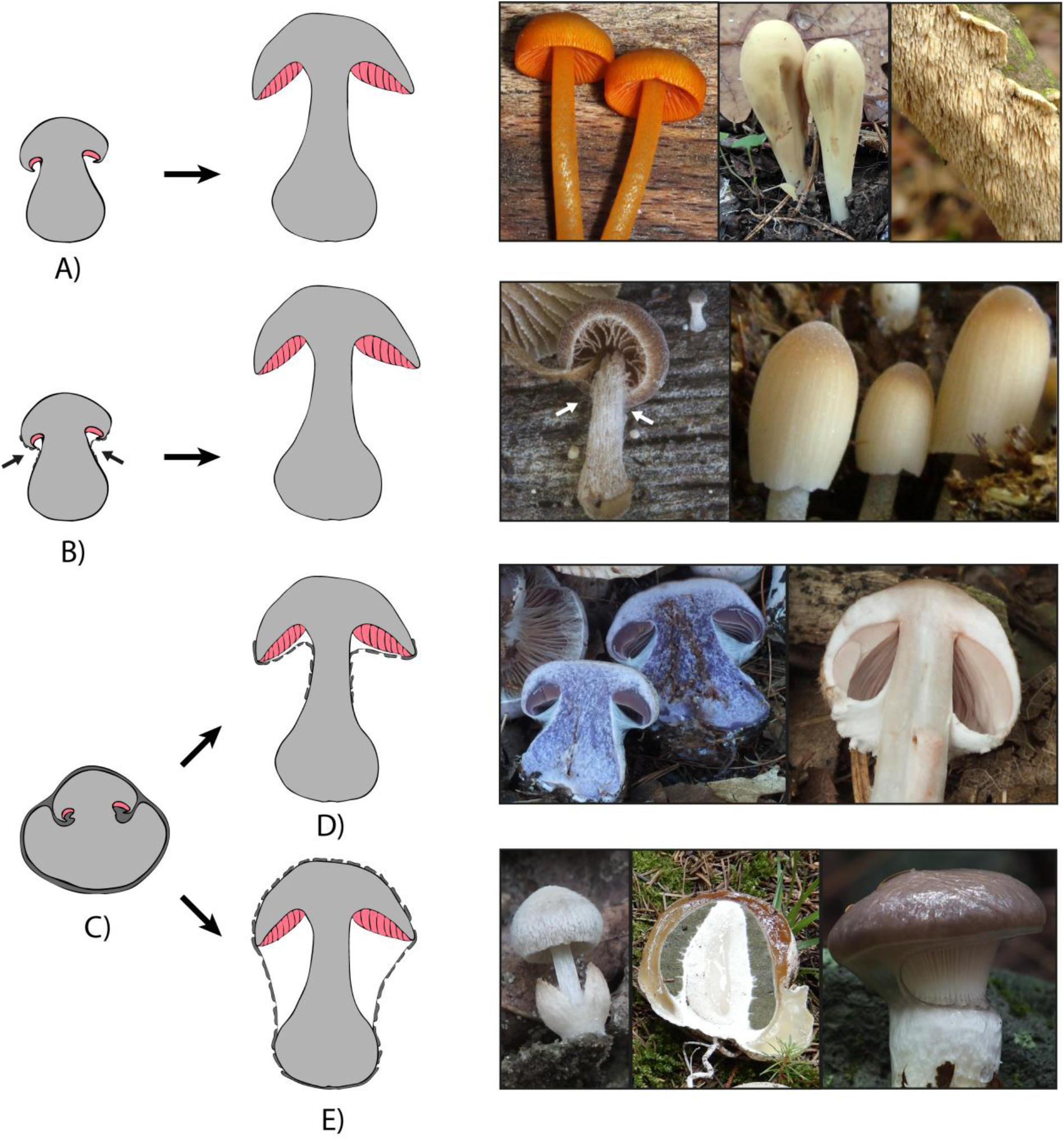
Developmental types in mushroom-forming fungi. Drawings depict primordial (left) and young fruiting bodies (right) of different developmental types. Magenta color shows the hymenial tissues. Dark grey color indicates tissues having role in the enclosure of the developing fruiting body A) Open development state. Images (left to right): Mycena leiana, Clavariadelphus pistillaris and Irpex lacteus B) Semi-enclosed development state. Note the faint tissue layer covering the hymenium of the primordium (arrowheads). Images (left to right): Ramicola sp. and Coprinellus congregatus. C) Enclosed development state; a robust tissue covers either the whole primordium or the hymenophore. D) and E) are subtypes of the enclosed development state showing partial and universal veils, respectively. D) Young fruiting body with partial veil Images (left to right): Cortinarius sp., Agaricus silvaticus. E) Young fruiting body with universal veil. Images (left to right): Volvariella sp., Phallus impudicus. Gomphidius glutinosus. Image courtesy: Alexey Sergeev, Judit Tóth Kőszeginé, László G. Nagy, Torda Varga.

To accurately define character states and thoroughly investigate the development of protecting tissue layers, we examined all 52 previous histological studies we could identify (table S1). From these studies, we made phylogenetically informed extrapolations to whole genera except for species with unique morphologies. In addition to plectological information, we gathered data on veil structures from the literature. In rare cases, we visually inspected images of young fruiting bodies and veil or tissue remnants on the cap and stipe. If assigning a state to species with confidence was not possible, we coded it as an ambiguous state. It is important to note that this character coding strategy lumps together multiple, traditionally recognized morphologies along the main criterion of exposure of hymenium (e.g., resupinate and coralloid forms in open development, or certain boletoid and agaricoid species in enclosed development).

To explore the robustness of the results to character state coding, we developed four alternative character state coding regimes (table S1). This was necessary as the extent of the hymenophore enclosure shows a continuum between fully open and enclosed. We broke up this continuum into discrete states to our best judgements and thoroughly examined if any bias could be introduced by the discretization of the character (see below). A character coding regime was created where a fourth character state was assigned to species with sequestrate or gasteroid fruiting bodies (referred to as 4ST1). We created further two modified versions of this four-state character coding. First, character states for certain ambiguous taxa were changed (4ST2): *Cribbea* spp. from state 0/3 to state 0, *Crinipellis* spp., *Lactarius* spp., *Marasmiellus* spp., *Tetrapyrgos* spp. from state 0/1 to state 0, *Deconica* spp., *Lactarius* spp. from state 0/2 to state 2, *Galerina* spp. from state 1/2 to state 2, *Pleuroflammula* spp. from state 1 to state 2, *Phaeocollybia* spp. from state 1/2 to state 0, some *Marasmius* spp. and *Mycena* spp. from state 1 to state 0 and *Naucoria* spp. from state 0/1 to state 2. Second, we re-coded all marasmoid fungi to state 0 (four states dataset 3, 4ST3) because certain histological studies (e.g., Clémencon, 2012; Reijnders, 1983) described only a faint and loose tissue layer between the cap and stipe at very early developmental stages. In the fourth alternative coding regime, to distinguish cyphelloid fungi from state 0, we produced a five-character state coding (5ST).

Finally, to examine the effects of multiple binary traits in one model, we created a binary coding by merging the semi-enclosed with the enclosed state of the 3ST coding regime into one state (2ST1). This appeared feasible because these two states behaved similarly in the trait dependent diversification analyses. In addition to this, we created a binary coding where semi-enclosed and open development were merged (2ST2) and where we randomly distributed the semi-enclosed state between species with enclosed or open development states (2ST3).

#### Partial and universal veil and hymenophore

We coded veil character states for each of the species in the phylogeny as two binary traits as follows (table S1). The partial veil (Clémencon, 2012) is defined as hyphal tissues that grow between the cap margin and the stipe and covers only the developing cap (including the hymenophore). The universal veil, on the other hand, covers the entire young fruiting body. In cases where a veil’s presence was not clear, we coded the species as ambiguous. In a few cases the nature of a veil was hard to define, therefore we created a veil coding where species with any of the veils were coded as state 1 and species without veils to state 0.

We included two additional morphological traits that could influence diversification rates: cap formation and increased hymenophore surface area. We obtained character coding for the cap from (Varga et al. 2019). For the hymenophore, we distinguished three character states based on the structural complexity of the hymenophore surface (table S1). Character state 0 was assigned to species with a smooth hymenophore. Character state 1 was assigned to species with weakly-structured hymenophore, which barely increases the hymenophore’s surface (e.g., veins, ridges, bumps). Character state 2 was assigned to species with complex hymenophores (e.g., gills, pores, teeth) (figure 2). To include the hymenophore into a multitrait binary model, we merged the smooth and weakly-structured states into state 0, and we assigned state 1 to species with complex hymenophore.

**Figure 2.**
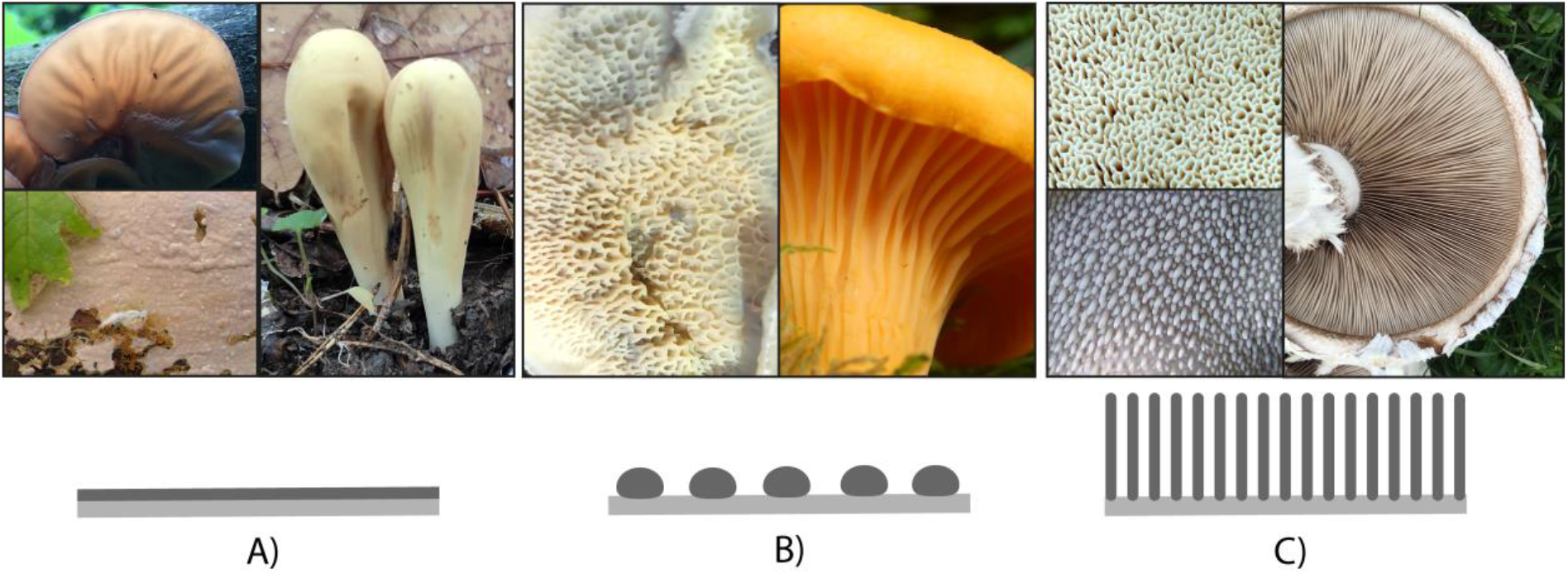
Three states/grades of hymenophore complexity distinguished in this study. A) Smooth hymenophore. Images: Auricularia auricula-judae (top), Clavariadelphus pistillaris (right) and Cylindrobasidium sp. (bottom). B) Weakly-structured hymenophore. Images: Phlebia tremellosa and Cantharellus cibarius. C) Complex hymenophore. Images: Bondarzewia montana (top), Nemecomyces mongolicus (right) Hydnum repandum (bottom). In schematic figures grey and dark grey denote supporting tissue (e.g., trama, subiculum) and sporogenous tissue (hymenium), respectively. Image courtesy: Judit Tóth Kőszeginé, Valéria Borsi, László G. Nagy, Torda Varga.

### Ancestral state reconstruction

Maximum parsimony (MP) based ancestral character state reconstruction (ASR) was performed using hsp_max_parsimony function from the castor v.1.5.5. R package (Louca & Doebeli, 2018) to calculate the number of origins of the different character states. We used a weighted transition cost matrix created from the transition rates inferred in the BayesTraits analysis (see below). First, we calculated the mean of the transition rates inferred through Bayesian and ML analysis in BayesTraits. Then we took the reciprocal of the values and shifted all values by the maximum of the reciprocal matrix to create a more significant gap between no cost (diagonal of the cost matrix) and the lowest cost transition. The hsp_max_parsimony function handles ambiguous character states as unknown states. To calculate the number of origins in ten chronograms of 5,284 species, we used a custom R function available at github.com/vtorda/ASR_analysis.

### Character state evolution

To infer macro-evolutionary transition rates, we used Maximum Likelihood (ML) and Markov Chain Monte Carlo (MCMC) approaches implemented in BayesTraits 2.0 Linux 64 Quad Precision alternative build (Meade & Pagel, 2016) and in diversitree 0.9-10 R (Fitzjohn, 2012). BayesTraits analyses were performed on 245 phylogenetic trees using the Multistate module of the program. We chose a gamma hyper-prior distribution for transition rates (table S2), which empirically fit best the data. This was determined based on preliminary analyses with uniform, exponential and gamma priors with and without a hyper-prior.

We observed high transition rate from semi-enclosed to open development which we hypothesised was caused by spuriously inferring an early gain of semi-enclosed, followed by frequent reversals to open development. To address this, we performed two additional tests. First, we examined whether constraining the stem nodes of 13 class- or order-level clades (Dacrymycetes, Cantharellales, Sebacinales, Auriculariales, Phallomycetidae, Trechisporales, Hymenochaetales, Boletales, Russulales+Polyporales, the hygrophoroid clade *sensu* Matheny (Matheny et al., 2006), Atheliaceae+Pterulaceae+Pleurotaceae, Physalacriaceae, agaricoid clade *sensu* Matheny et al. 2006) to open development state affects the transition rates. Second, we examined the contribution of individual clades to transition rates by setting state 1 or 01 of all species in a clade at a time to 0 and state 12 to 2. The rationale of this test was that a dramatic change in the transition rates relative to the original values, could mark a given clade as the main contributor to the global pattern. The following clades *sensu* Matheny et al. 2006 were examined with this procedure: Marasmioid clade, Tricholomatoid clade, Agaricoid clade, Psathyrellaceae, Boletales and Russulales (table S1).

All preliminary BayesTraits analyses (prior selection, constraining deep nodes, clade specific character state coding) were conducted with the following settings: 1,010,000 generations, 10,000 burn-in and sampling every 500^th^ generation. We observed that MCMC generally visited only ∼15 out of the 245 trees, which means 230 trees did not contribute toward our results (probably because Markov chains sampled trees in proportion to their likelihood). To overcome this, we forced the chain to spend 200,000 generations on each tree by the *EqualTrees* command and applying 100,000 burn-in and sampling every 500^th^ generation (altogether 49 million generations).

We used a less computationally demanding strategy for alternative character state coding regimes. In the case of 4ST1, 4ST2, and 4ST3 coding regimes, Markov chains were run for 10 million, while in the case of 5ST for 20 million generations with 10% burn-in and sampling every 500^th^ generation.

Marginal likelihoods were estimated by the stepping stone method (Meade & Pagel, 2016; Xie et al., 2011) using 50 stones with chain lengths of 5,000. Every analysis in BayesTraits was repeated three times to check the congruence of independent runs.

We performed model tests by comparing the unconstrained model and a nested model where certain constraints were made on the parameters. First, we tested if there is a tendency towards the evolution of any character states by constraining forward and reverse transition rates to be equal. This means one or three pair-wise constraints (q_01_ = q_10_, q_12_ = q_21_, q_02_ = q_20_) in case of binary or three state coding, respectively, and a constraint where all rates are equal (q_01_ = q_10_ = q_12_ = q_21_ = q_02_ = q_20_). To explore if a particular transition rate is supported by the data, we set the rate to zero (q_10_=0 or q_01_=0 or q_21_=0 or q_12_=0 or q_20_=0 or q_02_=0). Each of the constrained models mentioned above were compared to the best fit model using log-likelihood ratios (LR, ML analyses) or the log marginal likelihood ratio (Bayes factor, MCMC analyses). As a rule of thumb LR > 4 or Bayes factors > 10 was considered as significant support (Pagel, 1999).

Using ten chronograms, we also inferred transition rates and performed model testing under the multistate speciation and extinction (MuSSE, Fitzjohn, 2012) or the binary state speciation and extinction (BiSSE, Maddison et al., 2007) models for enclosed development 2ST and 3ST, increased hymenophore 3ST and 2ST, universal veil and partial veil traits. Significant differences among alternative models were determined by the likelihood ratio test (LRT) and Akaike information criterion scores (Fitzjohn, 2012; Meade & Pagel, 2016; Pagel, 1999), where p < 0.05 was considered to be significant. In case the enclosed development 2ST3 coding regime, we generated 100 perturbed traits by randomly distribute the semi-enclosed state to the two other states. Using this dataset and ten chronograms we performed 1,000 ML BiSSE analyses to infer transition, speciation and extinction rates.

### Trait-dependent diversification analyses

We used ten chronograms from our previous work (Varga et al., 2019) to analyze trait dependent diversification using the MuSSE or the BiSSE models implemented in diversitree v.09-10 R (Fitzjohn, 2012; Maddison et al., 2007). Transition, speciation, and extinction rates were inferred by using both ML and Bayesian MCMC. Starting points of ML searches were determined by the functions *starting*.*point*.*musse, starting*.*point*.*bisse*, and the analyses were corrected by state-specific sampling fractions (table S3) calculated by using our previous procedure, based on the number of species in Species Fungorum (Varga et al., 2019). Bayesian MCMC was performed using an exponential prior (defined by *1/(2r)*, where *r* is the character independent diversification rate) and Markov chains were run for 20,000 generations with 10% burn-in. The MCMC sampler’s step size was optimized after running 100 generations. Convergence of chains was inspected based on the variation of parameter values as a function of the number of generations.

We performed LRT to compare alternative models. We constrained state-specific speciation or extinction rates to be equal (λ_0_ = λ_1_ or λ_0_ = λ_2_ or λ_1_ = λ_2_ or μ_0_ = μ_1_ or μ_0_ = μ_2_ or μ_1_ = μ_2_) and performed LRT on the unconstrained model and the constrained models. We also tested whether a particular speciation or extinction rate is a significant component of the model by constraining it to zero (λ_0_ = 0 or λ_1_ = 0 or λ_1_ = 0 or μ_0_ = 0 or μ_1_ = 0 or μ_1_ = 0).

To analyze the effect of multiple binary traits on speciation and extinction rates in one model simultaneously, we used multitrait MuSSE model (Fitzjohn, 2012). The parameterization of the multitrait MuSSE model is analogous to that of a linear regression model. Consequently, an intercept (a “background rate”) and main effects (the effect of state 1 of any traits) are inferred. We analyzed two trait combinations: enclosed development – increased hymenophore surface area and enclosed development – cap. First, we compared the model where only the intercept was inferred (“depth” argument = c(0,0,0)) with the model where the main effect of the diversification was included (“depth” argument = c(1,1,0)) with performing LRT. Next, we carried out a Bayesian MCMC analysis using an exponential prior (defined by 1/(2r), where *r* is the character independent diversification rate), and Markov chains were run for 20,000 generations with 10% burn-in. We also examined the significance of the main effects of the binary traits by performing LRT on models where the effect of one of the traits was constrained to be 0 (λ_A_ = 0 or λ_B_ = 0). Finally, we compared the posterior distribution of parameter estimates of the multitrait MuSSE model and that of BiSSE models to examine how the speciation and extinction rates changed when analyzed within one model.

To rule out the possibility that the diversification rate pattern of enclosed development is driven by a trait which is not examined in this paper, we performed analyses by under the hidden state speciation and extinction (HiSSE) model (Beaulieu & O’Meara, 2016) implemented in RevBayes (Höhna et al., 2016). We defined a general HiSSE model where an observed binary and a hidden binary trait were included, and each of the four states can affect the diversification rate. We generated posterior samples by MCMC. Following the RevBayes manual (Höhna et al., 2019), we set a log-normal prior distribution for hidden speciation and extinction rate with a median of 1 and a standard deviation drawn from an exponential distribution with an empirical mean (0.587405). We specified a log-uniform distribution on the speciation and extinction rates of the observed trait between 10^−6^ and 10^2^. For observed and hidden transition rates, we defined an exponential prior with the mean of 10 transitions / total tree length. The sampling fraction parameter of the birth-death model was set to 0.148995, which was calculated based on known species numbers in the Species Fungorum database. We ran three independent chains for 4,500 generations each. Convergence was assessed by visually inspecting the saturation of likelihood and parameter values. We also evaluated geweke diagnostic plots (Geweke, 1992) implemented in the coda v.0.19-3 R (Plummer et al., 2006). We discarded samples prior to convergence and calculated effective sample sizes (ESS) for all parameters by the effectiveSize function.

## Results

### Enclosed development is the favored direction of evolution

We examined whether there is a trend towards enclosed development in the evolution of mushroom-forming fungi (Agaricomycetes) using comparative phylogenetic methods on previously published phylogenies of 5,284 species (Varga et al., 2019). MP ancestral state reconstructions on ten chronograms (table S4, figure S1) suggested that that the most recent common ancestor of the Agaricomycetes likely had open development. Semi-enclosed and enclosed development evolved 19-23 and 72-83 times, respectively, depending on the tree analyzed. Reversals to open development may have also happened 15-22 times. If we examined only internal nodes, 4-8, 11-14, and 25-31 transitions were inferred to open, semi-enclosed and enclosed development (figure 3), indicating that a significant proportion of transitions happened deeper in the tree. In line with these results, model inferences under maximum likelihood indicated significant asymmetries in transition rates among developmental types (figure 3). The highest average transition rates were inferred for the transition from the semi-enclosed to the enclosed state (q_12_); these rates were 8.6 - 21.0 times higher than the reverse rates (q_21_). Our results also suggest that the reversal from semi-enclosed to open development (q_10_) is frequent across the phylogeny. Model comparisons indicated that these asymmetric rate values were also significantly different from each other in all cases (LRT, p < 0.05, log Bayes Factor > 10; tables S2 and S5). Model testing also suggested that all transition rates were crucial parameters of the evolutionary model (i.e., significantly greater than zero, LRT, p < 0.05, Bayes factor > 10).

**Figure 3.**
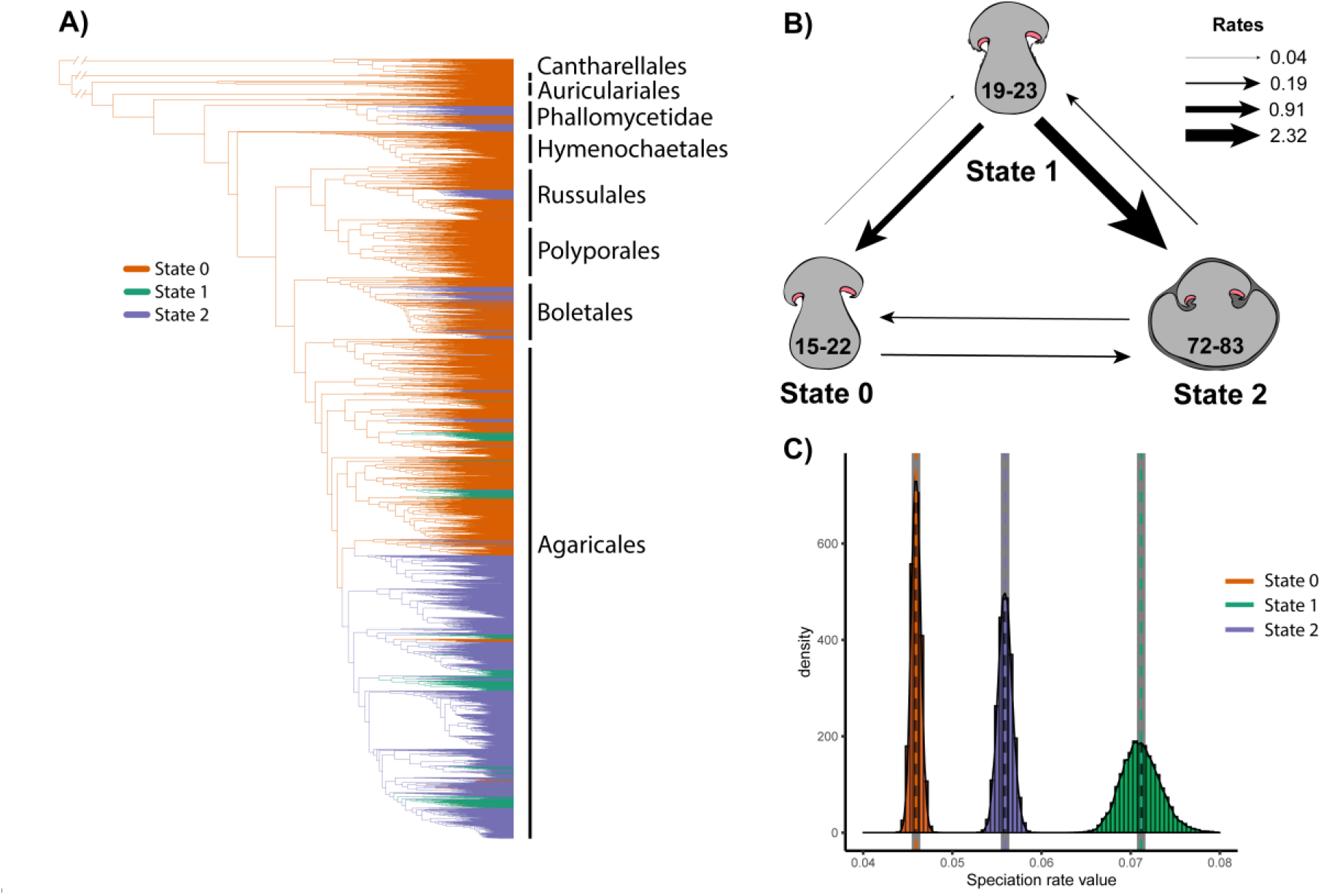
Macro-evolutionary patterns of enclosed development. A) Maximum parsimony ancestral state reconstruction of the 3ST coding regime. State 0 – Open development, State 1 - Semi-enclosed development, State 2 – Enclosed development. B) Evolutionary transitions between open (state 0) semi-enclosed (state 1) and enclosed (state 2) development. Number intervals on each schematic graphics of the states show the number of times a state evolved as inferred by maximum parsimony based on 10 trees. Arrows denote transition rates between states, their width is proportional to the mean transition rates inferred by Bayes Traits. C) Histograms show the posterior probability distribution of state-dependent speciation rates inferred by MuSSE.

Overall, these analyses suggest that enclosed development is a frequently-evolving and stable character state and its evolution is the preferred direction in mushroom-forming fungi. It may emerge either via a semi-enclosed intermediate (mean rate, q_01_ = 0.02 and q_12_ = 1.16) or could directly evolve from open development (mean rate, q_02_ = 0.10). Given that this developmental type provides the strongest physical protection from the environment of the three character states (but also requires the largest nutritional investment), it is conceivable that it confers a fitness advantage for mushroom-forming fungi, especially for those that produce above-ground fruiting bodies. On the other hand, the semi-enclosed state appears evolutionarily labile; once evolved, it either transforms into more persistent protective structures (enclosed state) or is lost rapidly (reversal to open), possibly due to the fugacious, incomplete protection it can provide to fruiting body initials.

Ancestral position of open development and convergent evolution of enclosed forms was also speculated in the Ascomycota. Phylogenetic studies of Pezizomycotina placed species with open fruiting bodies (apothecia) basally those with closed ones (perithecia) in more derived positions (Liu & Hall, 2004). In a lichen-forming ascomycetes group (Lecanoromycetes), several independent occurrences of enclosed (angiocarp) fruiting bodies were detected (Schmitt et al., 2009).

### Robustness to alternative character state codings

As character coding is always associated with a degree of subjectivity, we identified four potential major sources of subjectivity and addressed whether the above conclusions hold under alternative coding regimes. First, we tested if highly derived fruiting body morphologies (e.g., gasteroid or cyphelloid species) disproportionately contributed to the inferred patterns by recoding them as alternative character states (coding regimes 4ST1, 4ST2, 4ST3, 5ST). Second, we also tested if an early emergence of semi-enclosed development could cause spuriously high backwards transition rate to open development, or third, if certain major clades can individually drive the patterns observed above (figures S2-S3). Finally, given the difficulty of recognizing the semi-enclosed state, we addressed whether lumping it together with either of the other states, or randomly distributing it among them impacts our inferences (2ST1, 2ST2, 2ST3). Overall, we found that the transition rate pattern observed above (high q_10_ and q_12_ and low q_21_ and q_01_) was consistent across all alternative coding regimes (summarized in Supplementary Text S1, tables S2-S3). These findings indicate that our results are robust to character coding perturbations in the four most likely sources of subjectivity we identified.

### Enclosed development is associated with elevated species diversification rate

After we ascertained that the inferred transition rate patterns are robust, we evaluated the impact of enclosed development on speciation and extinction rates using state-dependent speciation and extinction (SSE) models. Species with semi-enclosed development have the highest net diversification rate (range of mean values across analyses: 6.5 × 10^−2^ – 8 × 10^−2^ events per million years), followed by species with enclosed development (5.5 × 10^−2^ – 6.1 × 10^−2^) and open development (4.6 × 10^−2^ – 5.2 × 10^−2^) (figure 3, Supplementary Table 3.), based on analyses of ten chronograms under the MuSSE model and ML or Bayesian methods. We found that speciation rate drove the differences in net diversification rates, because 26 out of 30 model tests showed significant differences in speciation rates (LRT, p < 0.05), but non-significant differences between any pair of extinction rates (Supplementary Table 5.). These results were robust to merging the semi-enclosed character state with either of the other two states and to randomly distributing it among other states (essentially reducing it to a BiSSE), implying that both semi-enclosed and enclosed development positively affect the diversification rate (Supplementary Text S1). We hypothesize that the elevated diversification rate stems from improved reproductive success conferred by the protection of fruiting body development, regardless of the complexity or the persistence of the given structure.

### Both partial and universal veils contribute to the diversification rate increase

Protection of fruiting body initials in species with enclosed development is provided by at least two morphological structures, the partial and universal veil, both of which could potentially drive increased diversification inferred above. The partial veil covers the hymenophore by stretching between the stem and the edge of the cap, whereas the universal veil envelopes the whole fruiting body when young. As the majority of species with enclosed or semi-enclosed development possess at least one kind of veil (figure 4), we attempted to dissect their contributions to diversification. The net diversification rate of species with universal or partial veils, respectively, was 1.23 and 1.33 times higher than that of species without either veil type (figure S4). As in the case of developmental types, diversification rate differences appear to be driven by differences in speciation rate, not extinction rate (LRT, p < 0.05). We found similar results when the universal and partial veil traits were combined into one trait (tables S3 and S5). These data suggest that both universal and partial veils contribute to increased diversification rates in species with (semi-)enclosed development. Although the exact ways in which veils increase fitness remain unknown at the moment, the upregulation of insecticidal and nematocidal toxin producing genes in veils suggest they may be involved in chemical and physical defense (Boulianne et al., 2000; Sabotič et al., 2011).

**Figure 4.**
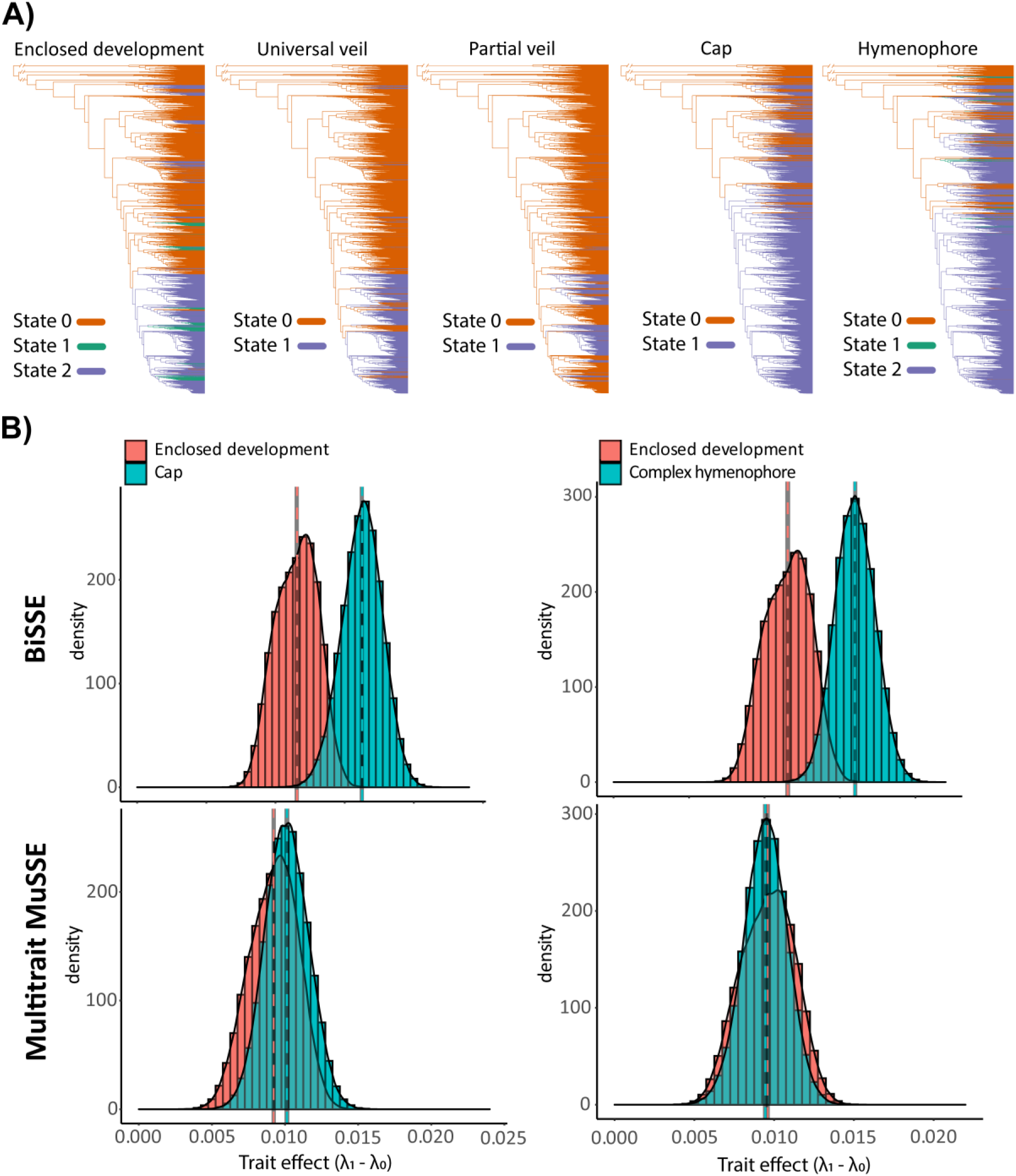
A) Maximum parsimony ancestral reconstruction of five morphological characters examined in this study. Enclosed development trait (open development - state 0, semi-enclosed development – state 1, enclosed development – state 2). Universal veil trait (absence – state 0, presence – state 1). Partial veil trait (absence – state 0, presence – state 1). Cap trait (absence – state 0, presence – state 1). Hymenophore complexity trait (Smooth hymenophore - state 0, weakly-structured hymenophore – state 1, complex hymenophore – state 2). B) speciation rate effects (λ1-λ0) inferred by BiSSE (upper row) and multitrait MuSSE (bottom row) analyses. Histograms on the left show the comparison of the enclosed development and the cap traits. Histograms on the right show the comparison of the enclosed development and the hymenophore traits.

### The impact of enclosed development on diversification is independent of other observed or unobserved traits

Diversification rate differences can be driven by single or by interactions between multiple traits (Rabosky & Goldberg, 2015). We therefore tested whether the observed impact of developmental type on diversification rate could have been influenced by phylogenetically co-distributed characters (figure 4). We first examined the simultaneous effect of enclosed development and other morphological traits (morphological complexity of the hymenophore and the presence of a cap) in a multitrait speciation and extinction model (multitrait MuSSE). This model allowed us to decipher the background and the individual (“main trait”) effects of binary traits on diversification rate, and thus to separately evaluate the contribution of each trait to diversification rate changes (Fitzjohn, 2012).

In the multitrait MuSSE framework the models with main trait effects were superior over the model with only the “background” effect (LRT, p < 0.05). We found that enclosed development, the hymenophore and the cap were all significant components of the model with main trait effects, because the log-likelihoods of the unconstrained models were significantly higher than that of models where the effect of one or the other trait was constrained to zero (LRT, p<0.05). We found that the speciation rate differences under two states (λ_1_-λ_0_, called ‘trait effect’) in the multitrait analyses were lower than those in the BiSSE analyses (figure 4). This indicates that speciation rate differences inferred under BiSSE are, to an extent, arise from the interaction of two traits. However, the speciation rates of lineages with any of the traits remained significantly higher (λ_1_-λ_0_>0, LRT, p < 0.05, figure 4) than that of clades without the trait, indicating a robust and independent positive impact on diversification by each of the three traits (enclosed development, hymenophore, or the presence of a cap). This suggests that the increased diversification of species with enclosed development is independent of both the complexity of the hymenophore or the presence of a cap.

To address the possibility that other unobserved or hidden traits drove the observed patterns, we performed a hidden state speciation and extinction (HiSSE) analysis. We found that the speciation rate of species with enclosed development was higher than that of non-enclosed (figure S5), even in the presence of a hidden trait in the HiSSE model, suggesting that the diversification rate patterns we identified are indeed attributable to innovations in developmental mode.

### Hymenophore complexity alone also impacts diversification

We were also curious whether hymenophore complexity alone influenced species diversification (the impact of the cap has been examined before, see Varga et al 2019). MP ancestral state reconstruction suggested that the most common ancestor of Agaricomycetes was a mushroom with smooth hymenophore and that weakly-structured and complex hymenophores evolved 41 - 46 and 74 - 91 times, respectively (table S4). BayesTraits and MuSSE analyses showed that the transition rate from weakly-structured towards complex hymenophore (q_12_) was 54.3 - 61.5 times higher than in the reverse direction (q_21_) and significantly ‘non-equal’ (figure 5, LRT, p < 0.05, log Bayes factor > 10). We also found that the transition rate from weakly-structured hymenophore towards smooth hymenophore was 13.4 – 17.3 times higher than in the reverse direction (LRT, p < 0.05, log Bayes factor > 10), suggesting several reversals of weakly-structured hymenophores to smooth ones. This implies that complex hymenophores with increased surface area are favored during the evolution of mushroom-forming fungi.

**Figure 5.**
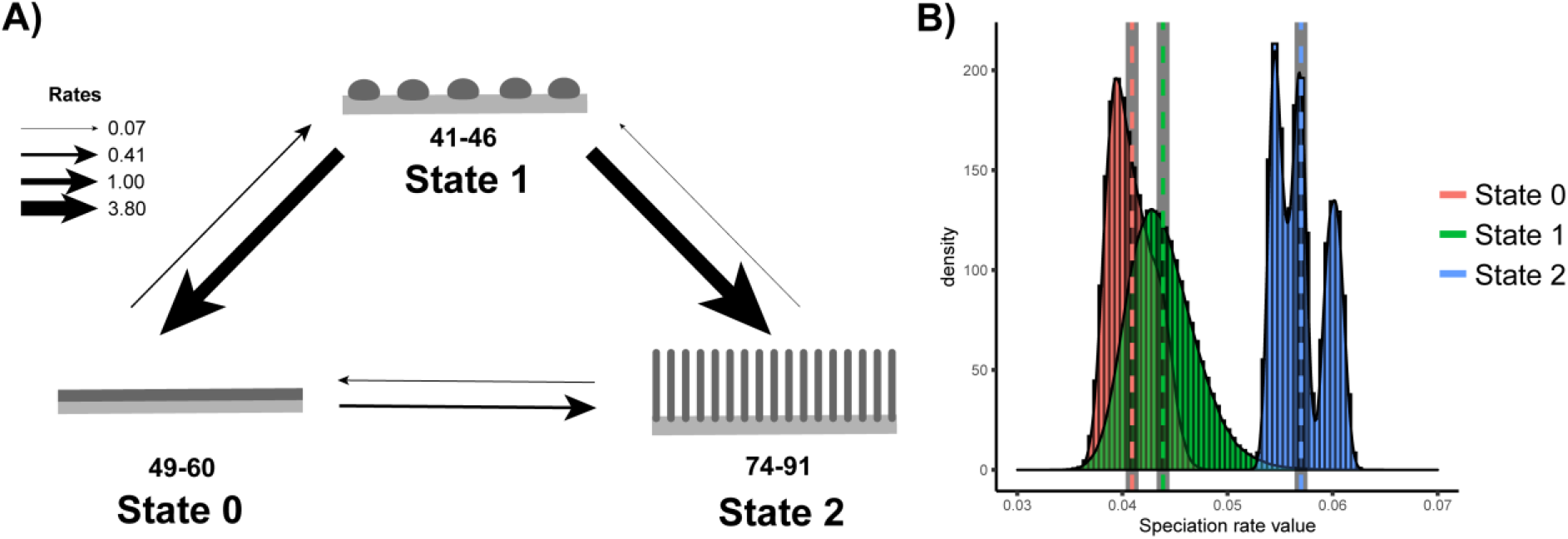
Macro-evolutionary patterns of hymenophore complexity. A) Transition rates between the three character states (state 0 – smooth hymenophore, state 1 – weakly-structured hymenophore, state 2 – complex hymenophore), inferred by Bayes Traits. The intervals below the schematic graphics represent the number of times a state evolved according to maximum parsimony ancestral state reconstruction. The width of the arrows is proportional to the transition rates. B) Histograms depicting the state dependent diversification rates of the three character states inferred by MuSSE.

We found that the diversification rate of species with complex hymenophore was significantly higher than that of species with smooth or weakly-structured hymenophore (LRT, p < 0.05), while diversification rates of species with the latter two did not differ significantly (figure 5, LRT, p >0.05). These results suggest that only well-developed gills, pores, and teeth can positively affect diversification of mushroom-forming fungi, whereas weakly-structured hymenophores (bumps, ridges, veins) do not and they revert frequently to smooth surfaces.

## Conclusions

The evolutionary success of species is strongly connected to their reproductive efficiency. Accordingly, the impact of innovations on reproductive ability influence which morphologies, behaviors or other traits reach high equilibrium frequencies or go extinct in their clades. In the context of fungi, traits related to spore production, dispersal and germination are among the primary determinants of reproductive success in terrestrial habitats (Aguilar-Trigueros et al., 2019; Halbwachs et al., 2015; Hibbett & Binder, 2002; James, 2015; Norros et al., 2014; Peay et al., 2012). Such traits should, thus, drive morphological evolution in sexual fruiting bodies and should impact lineage diversification. Despite this clear prediction, what adaptations fruiting bodies evolved for increasing spore dispersal efficiency are hardly known and studies addressing the correlations between morphogenetic traits and species diversification are at paucity.

In this study, we provide evidence that morphological innovations pertaining to the efficiency of spore production show considerable asymmetry in their evolution and their evolution is associated with increased diversification rates (i.e., may be key innovations) in mushroom-forming fungi. These include enclosed development, in which fruiting body initials are ensheathed by veil tissues, providing protection to the fruiting body initial, in a manner analogous to the internally nursed embryos of viviparous animals and plants (though its important to note that the fruiting body initial serves a different purpose from the plant/animal embryo). In analogy, viviparity spurred lineage diversification in squamates and cyprinodontiform fishes (Helmstetter et al., 2016; Pyron & Burbrink, 2014). Our analyses provided clear support for convergent origins of and asymmetrical evolution favoring enclosed development and a correlation with increased lineage diversification rates. These results were robust to model and method choice, alternative coding regimes and not affected by character states at basal nodes or in any major clades.

Protecting fruiting body initials is of prime importance as these contain the developing hymenium, on which basidia and spores are born.Albeit fruiting bodies generally quickly complete their developmental program and sporulate (though in some species the process can take weeks), several factors can compromise development (desiccation, predators, infections, rain, other physical damages) and consequently impede sporulation. It has been shown that secondary metabolites, peptides, proteins (e.g., galectins) against bacteria or fungivorous animals (mammals, arthropods, nematodes) are produced by tissues that ensheath fruiting body initials (Bleuler-Martínez et al., 2011; Boulianne et al., 2000; Jaeger & Spiteller, 2010; Künzler, 2018; Sabotič et al., 2011, 2016). Enclosed development might also help phasing the growth of fruiting bodies by providing a sheltered environment for cell differentiation, after which rapid growth by cell expansion (Kües, 2000) lifts the hymenophore quickly above ground. This might be advantageous in terrestrial habitats, where developing at ground level and lifting the cap above ground reduced evaporation and potentially allows the development of larger fruiting bodies, which increase spore quantity and release height, two critical factors in dispersal (Norros et al., 2014).

At large evolutionary scales such as the one examined in this paper, causes of diversification rate differences may easily be distributed among a nested set of phenotypic innovations or phylogenetically co-distributed traits (Donoghue, 2005). To address this possibility, we examined two velar structures, which alone or in combination provide the physical barriers to the environment in most species with enclosed development, as well as alternative, phylogenetically nested or unknown traits. We found that both universal and partial veil and their combination associate with differences in diversification rates, suggesting that enclosure of the hymenophore by either veil type is sufficient for diversification rates to increase. We further examined the effects of two independent, but conceivably adaptive traits, the presence of a cap and structured hymenophore surfaces. Multitrait BISSE models, which test the combined effects of multiple traits on diversification in a single analysis, provided evidence that, albeit both traits influence diversification rates (see also Varga et al., 2019), their effects are independent from that of enclosed development. Complex hymenophore itself was, independently of enclosed development, associated with higher diversification rates relative to simpler morphologies (smooth or weakly-structured hymenophores), suggesting that hymenophoral complexity is adaptive in mushroom-forming fungi, possibly by allowing the production of more propagules per unit biomass (Iapichino et al., 2021), variations in gill positioning (Fischer & Money, 2010), protection against predators (Nakamori & Suzuki, 2007), keeping high humidity (Halbwachs & Bässler, 2015) or producing local winds that help spores dispersal. Finally, hidden state speciation extinction analyses (Beaulieu & O’Meara, 2016) excluded the possibility that phylogenetically co-distributed and unobserved traits, rather than enclosed development is the main driver of diversification rate differences in mushroom-forming fungi.

Overall, these results give us confidence that the observed effect of enclosed development on diversification rates is robust to methods, dataset or other candidate traits which we tested. However, these analyses identified several traits which independently are associated with increases in diversification rates, indicating that beyond developmental types, the extant diversity of Agaricomycetes has probably been influenced by a complex interplay between multiple fruiting body and, possibly also nutritional innovations and that phylogenetically nested sets of these may underlie the radiation and evolutionary success of mushroom-forming fungi, one of the most diverse, important and spectacular components of the ecosystem.

## Acknowledgements

The authors acknowledge support by the “Momentum” program of the Hungarian Academy of Sciences (contract No. LP2019-13/2019 to L.G.N.) and the European Research Council (grant no. 758161 to L.G.N.). We are thankful to Valéria Borsi, Alexey Sergeev and Judit Tóth Kőszeginé for providing us the images of *Cantharellus cibarius, Ramicola* sp., as well as *Phallus impudicus* and *Hydnum repandum*, respectively.

## Descriptions of supplementary materials

Supplementary Text S1: Detailed description of the analyses of alternative coding regimes of enclosed development

Supplementary Table S1. Character coding data table and a summary of species and genera on which histological studies have been analyzed.

Supplementary Table S2. Parameters and model tests of the BayesTraits analyses.

Supplementary Table S3. State-specific sampling fractions and inferred parameters of the state-specific speciation and extinction (SSE) analyses.

Supplementary Table S4. Cost matrices used in maximum parsimony ancestral state reconstruction analyses and the number of transformations between states of enclosed development and hymenophore.

Supplementary Table S5. Model tests performed on state-specific speciation and extinction (SSE) analyses.

Supplementary Figure S1. A plot of maximum parsimony ancestral states of enclosed development on a randomly chosen tree (#8) from Varga et al. 2019. Green – open development, red – semi-enclosed development, blue – enclosed development.

Supplementary Figure S2. Visual representation of the BayesTraits analyses of enclosed development, where the stem node of 13 clades was constrained to state 0. A) The phylogenetic tree of Agaricomycetes with 13 clades of which stem node was constrained. B) Histograms of transition rates of the default analyses (red) and the analyses where the stem nodes were constrained to 0 (green).

Supplementary Figure S3. Density plots of transition rates between states of enclosed development inferred under alternative and default (3ST) character coding regimes. Alternative coding regimes were created by setting state 1 or 01 of all species in a clade at a time to 0 and state 12 to 2. Results are shown for six clades tested this way. Below each plot, a table showing mean and standard deviation of parameter estimates is given.

Supplementary Figure S4. Histograms depicting state-specific transition rates of the universal veil (A) and partial veil traits (B). The state-specific transition rates were inferred by Bayesian MuSSE analyses using ten chronograms of Agaricomycetes from Varga et al. 2019. lambda1 is the speciation rate of lineages with a universal (A) or partial veil (B) and lambda0 is the speciation rate of lineages without a universal (A) or partial veil (B).

Supplementary Figure S5. Comparative boxplots of open and enclosed development-specific speciation rates inferred by hidden state speciation and extinction (HiSSE) analysis. We compared the inferred speciation rate of lineages with enclosed (green) and open (red) development within each analysis of the ten randomly chosen chronograms.

## Notes

### Competing Interest Statement

The authors have declared no competing interest.

## References

Aguilar-Trigueros, C. A., Hempel, S., Powell, J. R., Cornwell, W. K., & Rillig, M. C. (2019). Bridging reproductive and microbial ecology: a case study in arbuscular mycorrhizal fungi. ISME Journal, 13(4), 873–884. https://doi.org/10.1038/s41396-018-0314-7

Beaulieu, J. M., & O’Meara, B. C. (2016). Detecting hidden diversification shifts in models of trait-dependent speciation and extinction. Systematic Biology, 65(4), 583–601. https://doi.org/10.1093/sysbio/syw022

Blackburn, D. G. (1999). Viviparity and oviparity: Evolution and reproductive strategies. Encyclopedia of Reproduction, 4(JANUARY 1999), 994–1003.

Blackburn, D. G. (2015). Evolution of vertebrate viviparity and specializations for fetal nutrition: A quantitative and qualitative analysis. Journal of Morphology, 276(8), 961– 990. https://doi.org/10.1002/jmor.20272

Bleuler-Martínez, S., Butschi, A., Garbani, M., WÎlti, M. A., Wohlschlager, T., Potthoff, E., Sabotia, J., Pohleven, J., Lüthy, P., Hengartner, M. O., Aebi, M., Künzler, M., Wílti, M. A., Wohlschlager, T., Potthoff, E., Sabotia, J., Pohleven, J., Lüthy, P., Hengartner, M. O., … Künzler, M. (2011). A lectin-mediated resistance of higher fungi against predators and parasites. Molecular Ecology, 20(14), 3056–3070. https://doi.org/10.1111/j.1365-294X.2011.05093.x

Boulianne, R. P., Liu, Y., Lu, B. C., Kües, U., & Aebi, M. (2000). Fruiting body development in Coprinus cinereus: regulated expression of two galectins secreted by a non-classical pathway. Microbiology, 146(8), 1841–1853. https://doi.org/10.1099/00221287-146-8-1841

Braga, G. U. L., Rangel, D. E. N., Fernandes, É. K. K., Flint, S. D., & Roberts, D. W. (2015). Molecular and physiological effects of environmental UV radiation on fungal conidia. Current Genetics, 61(3), 405–425. https://doi.org/10.1007/s00294-015-0483-0

Clémencon, H. (2012). Cytology and Plectology of the Hymenomycetes (2nd ed.). Gebr. Borntraeger Verlagsbuchhandlung.

Donoghue, M. J. (2005). Key innovations, convergence, and success: macroevolutionary lessons from plant phylogeny. Paleobiology, 31(2), 77–93. https://doi.org/10.1666/0094-8373(2005)031[0077:kicasm]2.0.co;2

Dressaire, E., Yamada, L., Song, B., & Roper, M. (2016). Mushrooms use convectively created airflows to disperse their spores. Proceedings of the National Academy of Sciences of the United States of America, 113(11), 2833–2838. https://doi.org/10.1073/pnas.1509612113

Fischer, M. W. F., & Money, N. P. (2010). Why mushrooms form gills: efficiency of the lamellate morphology. Fungal Biology, 114(1), 57–63. https://doi.org/10.1016/j.mycres.2009.10.006

Fitzjohn, R. G. (2012). Diversitree: Comparative phylogenetic analyses of diversification in R. Methods in Ecology and Evolution, 3(6), 1084–1092. https://doi.org/10.1111/j.2041-210X.2012.00234.x

Geweke, J. (1992). Evaluating the accuracy of sampling-based approaches to the calculation of posterior moments. Bayesian Statistics 4, 169–193. https://doi.org/1176289

Goldberg, R. B., de Paiva, G., & Yadegari, R. (1994). Plant Embryogenesis: Zygote to Seed. Science, 266(5185), 605–614. https://doi.org/10.1126/science.266.5185.605

Halbwachs, H., & Bässler, C. (2015). Gone with the wind – a review on basidiospores of lamellate agarics. Mycosphere, 6(1), 78–112. https://doi.org/10.5943/mycosphere/6/1/10

Halbwachs, H., Brandl, R., & Bässler, C. (2015). Spore wall traits of ectomycorrhizal and saprotrophic agarics may mirror their distinct lifestyles. Fungal Ecology, 17, 197–204. https://doi.org/10.1016/j.funeco.2014.10.003

Helmstetter, A. J., Papadopulos, A. S. T., Igea, J., Van Dooren, T. J. M., Leroi, A. M., & Savolainen, V. (2016). Viviparity stimulates diversification in an order of fish. Nature Communications, 7, 11271. https://doi.org/10.1038/ncomms11271

Hibbett, D. S. (2004). Trends in Morphological Evolution in Homobasidiomycetes Inferred Using Maximum Likelihood: A Comparison of Binary and Multistate Approaches. Systematic Biology, 53(6), 889–903. https://doi.org/10.1080/10635150490522610

Hibbett, D. S., & Binder, M. (2002). Evolution of complex fruiting-body morphologies in homobasidiomycetes. Proceedings of the Royal Society B: Biological Sciences, 49(191), 1963–1969. https://doi.org/10.1098/rspb.2002.2123

Höhna, S., Landis, M. J., Heath, T. A., Boussau, B., Lartillot, N., Moore, B. R., Huelsenbeck, J. P., & Ronquist, F. (2016). RevBayes: Bayesian Phylogenetic Inference Using Graphical Models and an Interactive Model-Specification Language. Systematic Biology, 65(4), 726–736. https://doi.org/10.1093/sysbio/syw021

Höhna, S., Ronquist, F., Landis, M. J., Boussau, B., Heath, T. A., Lartillot, N., Pett, W., Freyman, W. A., & Huelsenbeck, J. P. (2019). RevBayes: Bayesian phylogenetic inference using probabilistic graphical models and an interpreted language. https://Revbayes.Github.Io/.

Iapichino, M., Wang, Y.-W., Gentry, S., Pringle, A., & Seminara, A. (2021). A precise relationship among Buller’s drop, ballistospore, and gill morphologies enables maximum packing of spores within gilled mushrooms. Mycologia, 00(00), 1–12. https://doi.org/10.1080/00275514.2020.1823175

Jaeger, R. J. R., & Spiteller, P. (2010). Mycenaaurin A, an antibacterial polyene pigment from the fruiting bodies of mycena aurantiomarginata. Journal of Natural Products, 73(8), 1350–1354. https://doi.org/10.1021/np100155z

James, T. Y. (2015). Why mushrooms have evolved to be so promiscuous: Insights from evolutionary and ecological patterns. Fungal Biology Reviews, 29(3–4), 167–178. https://doi.org/10.1016/j.fbr.2015.10.002

Kües, U. (2000). Life history and developmental processes in the basidiomycete Coprinus cinereus. Microbiology and Molecular Biology Reviews?: MMBR59-, 64(2), 316–353. https://doi.org/10.1128/MMBR.64.2.316-353.2000

Künzler, M. (2018). How fungi defend themselves against microbial competitors and animal predators. PLOS Pathogens, 14(9), e1007184. https://doi.org/10.1371/journal.ppat.1007184

Liu, Y. J., & Hall, B. D. (2004). Body plan evolution of ascomycetes, as inferred from an RNA polymerase II phylogeny. Proceedings of the National Academy of Sciences of the United States of America, 101(13), 4507–4512. https://doi.org/10.1073/pnas.0400938101

Louca, S., & Doebeli, M. (2018). Efficient comparative phylogenetics on large trees. Bioinformatics, 34(6), 1053–1055. https://doi.org/10.1093/bioinformatics/btx701

Maddison, W. P., Midford, P. E., & Otto, S. P. (2007). Estimating a binary character’s effect on speciation and extinction. Systematic Biology, 56(5), 701–710. https://doi.org/10.1080/10635150701607033

Matheny, P. B., Curtis, J. M., Hofstetter, V., Aime, M. C., Moncalvo, J.-M. J.-M. M., Ge, Z.-W. Z.-W. W., Yang, Z. L. Z.-L., Slot, J. C., Ammirati, J. F., Baroni, T. J., Bougher, N. L., Hughes, K. W., Lodge, D. J., Kerrigan, R. W., Seidl, M. T., Aanen, D. K., DeNitis, M., Daniele, G. M., Desjardin, D. E., … Hibbett, D. S. (2006). Major clades of Agaricales: A multilocus phylogenetic overview. Mycologia, 98(6), 982–995. https://doi.org/10.3852/mycologia.98.6.982

Meade, A., & Pagel, M. (2016). BayesTraits V3. November, 81. http://www.evolution.rdg.ac.uk/BayesTraitsV3/Files/BayesTraitsV3.Manual.pdf

Nakamori, T., & Suzuki, A. (2007). Defensive role of cystidia against Collembola in the basidiomycetes Russula bella and Strobilurus ohshimae. Mycological Research, 111(11), 1345–1351. https://doi.org/10.1016/j.mycres.2007.08.013

Norros, V., Rannik, Ü., Hussein, T., Petäjä, T., Vesala, T., & Ovaskainen, O. (2014). Do small spores disperse further than large spores? Ecology, 95(6), 1612–1621. https://doi.org/10.1890/13-0877.1

Pagel, M. (1999). Inferring the historical patterns of biological evolution. Nature, 401(6756), 877–884. https://doi.org/10.1038/44766

Peay, K. G., Schubert, M. G., Nguyen, N. H., & Bruns, T. D. (2012). Measuring ectomycorrhizal fungal dispersal: macroecological patterns driven by microscopic propagules. Molecular Ecology, 21(16), 4122–4136. https://doi.org/10.1111/j.1365-294X.2012.05666.x

Plummer, M., Best, N., Cowles, K., & Vines, K. (2006). CODA: convergence diagnosis and output analysis for MCMC. R News, 6(March), 7–11. https://doi.org/10.1159/000323281

Pyron, R. A., & Burbrink, F. T. (2014). Early origin of viviparity and multiple reversions to oviparity in squamate reptiles. Ecology Letters, 17(1), 13–21. https://doi.org/10.1111/ele.12168

Rabosky, D. L., & Goldberg, E. E. (2015). Model inadequacy and mistaken inferences of trait-dependent speciation. Systematic Biology, 64(2), 340–355. https://doi.org/10.1093/sysbio/syu131

Reijnders, A. F. M. (1983). Supplementary notes on basidiocarp ontogeny in agarics. Persoonia, 12(1), 1–20.

Roberts, R. M., Green, J. A., & Schulz, L. C. (2016). The evolution of the placenta. Reproduction, 152(5), R179–R189. https://doi.org/10.1530/REP-16-0325

Sabotič, J., Kilaru, S., Budič, M., Gašparič, M. B., Gruden, K., Bailey, A. M., Foster, G. D., & Kos, J. (2011). Protease inhibitors clitocypin and macrocypin are differentially expressed within basidiomycete fruiting bodies. Biochimie, 93(10), 1685–1693. https://doi.org/10.1016/j.biochi.2011.05.034

Sabotič, J., Ohm, R. A., & Künzler, M. (2016). Entomotoxic and nematotoxic lectins and protease inhibitors from fungal fruiting bodies. Applied Microbiology and Biotechnology, 100(1), 91–111. https://doi.org/10.1007/s00253-015-7075-2

Sánchez-García, M., Ryberg, M., Khan, F. K., Varga, T., Nagy, L. G., & Hibbett, D. S. (2020). Fruiting body form, not nutritional mode, is the major driver of diversification in mushroom-forming fungi. Proceedings of the National Academy of Sciences, 117(51), 32528–32534. https://doi.org/10.1073/pnas.1922539117

Schmitt, I., Prado R. del, Grube, M., & Lumbsch, H. T. (2009). Repeated evolution of closed fruiting bodies is linked to ascoma development in the largest group of lichenized fungi (Lecanoromycetes, Ascomycota). Molecular Phylogenetics and Evolution, 52(1), 34–44. https://doi.org/10.1016/j.ympev.2009.03.017

Vamosi, J. C., Magallón, S., Mayrose, I., Otto, S. P., & Sauquet, H. (2018). Macroevolutionary Patterns of Flowering Plant Speciation and Extinction. Annual Review of Plant Biology, 69(February), 685–706. https://doi.org/10.1146/annurev-arplant-042817-040348

Varga, T., Krizsán, K., Földi, C., Dima, B., Sánchez-García, M., Sánchez-Ramírez, S., Szöllősi, G. J. G. J., Szarkándi, J. G. J. G., Papp, V., Albert, L., Andreopoulos, W., Angelini, C., Antonín, V., Barry, K. W. K. W., Bougher, N. L. N. L., Buchanan, P., Buyck, B., Bense, V., Catcheside, P., … Nagy, L. G. L. G. (2019). Megaphylogeny resolves global patterns of mushroom evolution. Nature Ecology & Evolution, 3(4), 668–678. https://doi.org/10.1038/s41559-019-0834-1

Westoby, M., & Rice, B. (1982). EVOLUTION OF THE SEED PLANTS AND INCLUSIVE FITNESS OF PLANT TISSUES. Evolution, 36(4), 713–724. https://doi.org/10.1111/j.1558-5646.1982.tb05437.x

Xie, W., Lewis, P. O., Fan, Y., Kuo, L., & Chen, M. H. (2011). Improving marginal likelihood estimation for bayesian phylogenetic model selection. Systematic Biology, 60(2), 150– 160. https://doi.org/10.1093/sysbio/syq085

