## Supplementary figures and images for "Developmental innovations promote species diversification in mushroom-forming fungi"

### figure S1

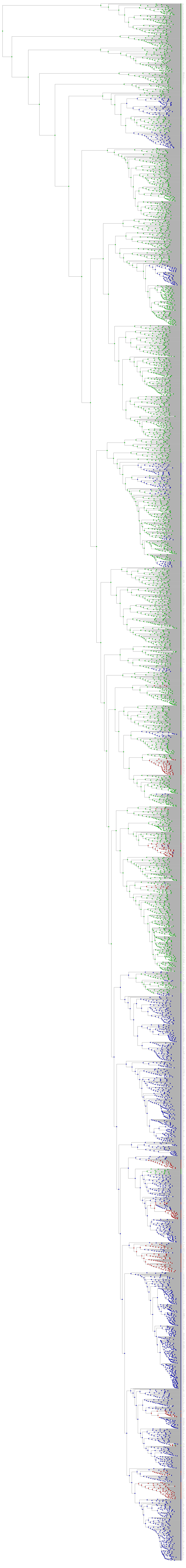

### figure S2

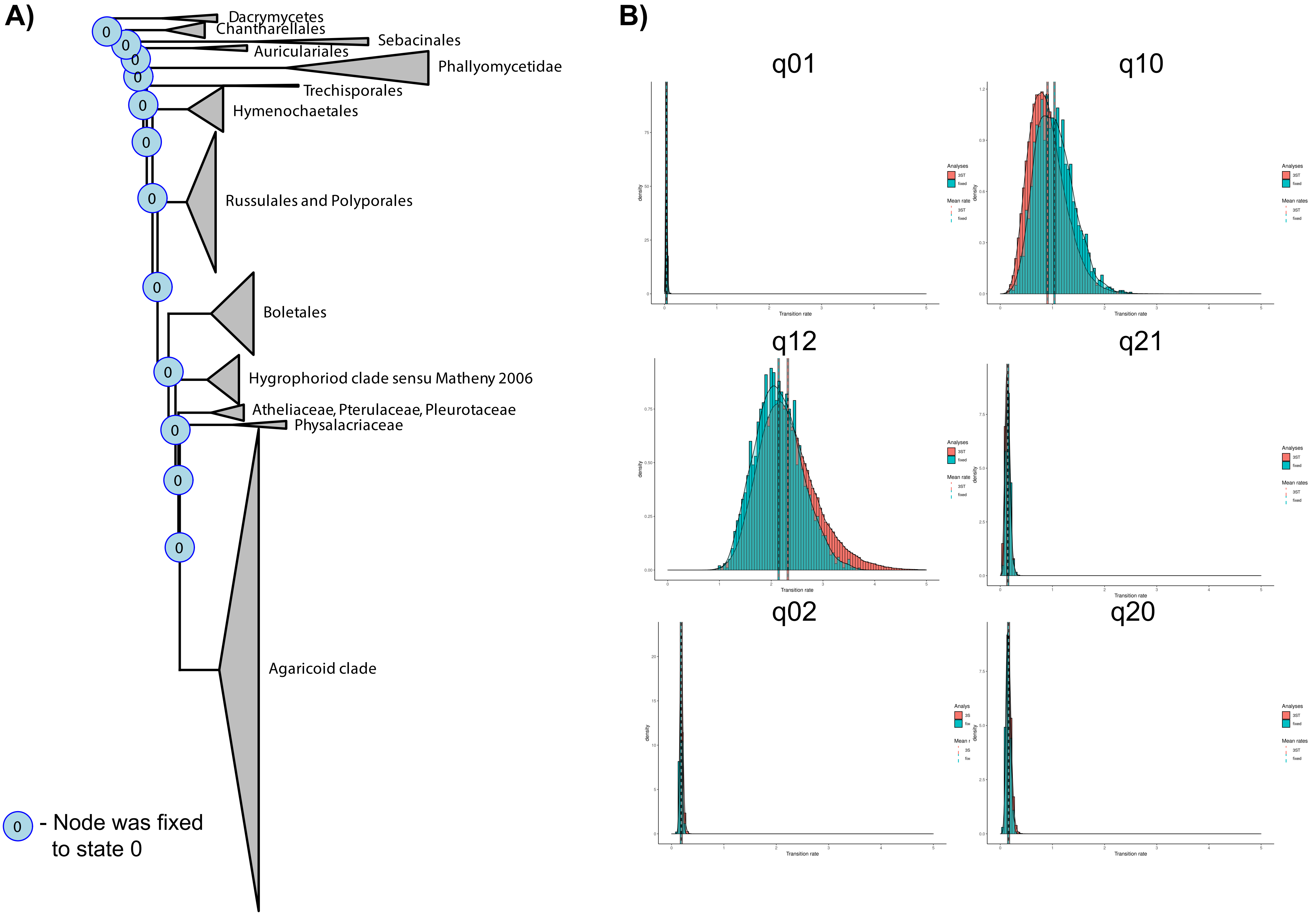

### figure S4

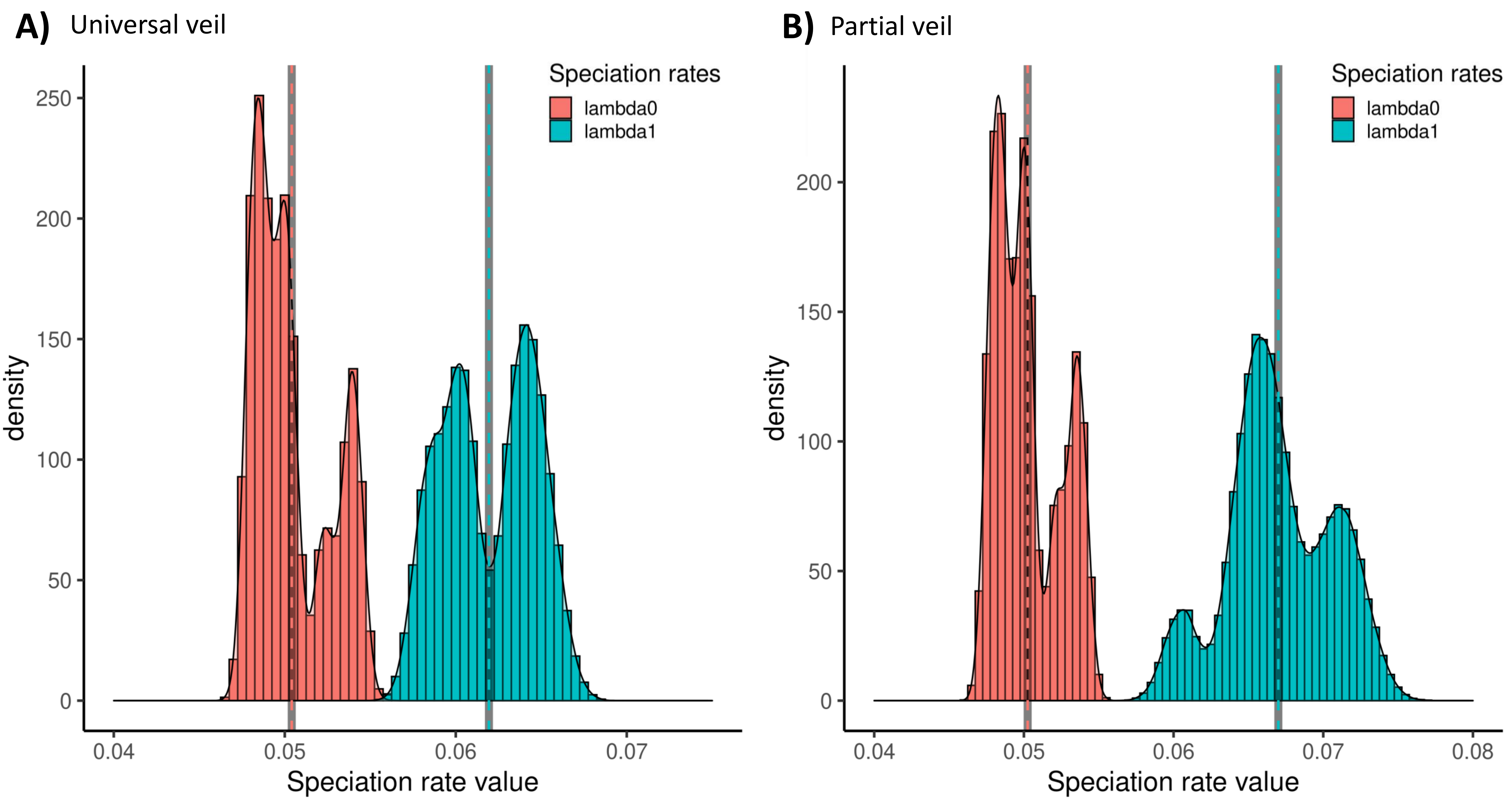

### figure S5

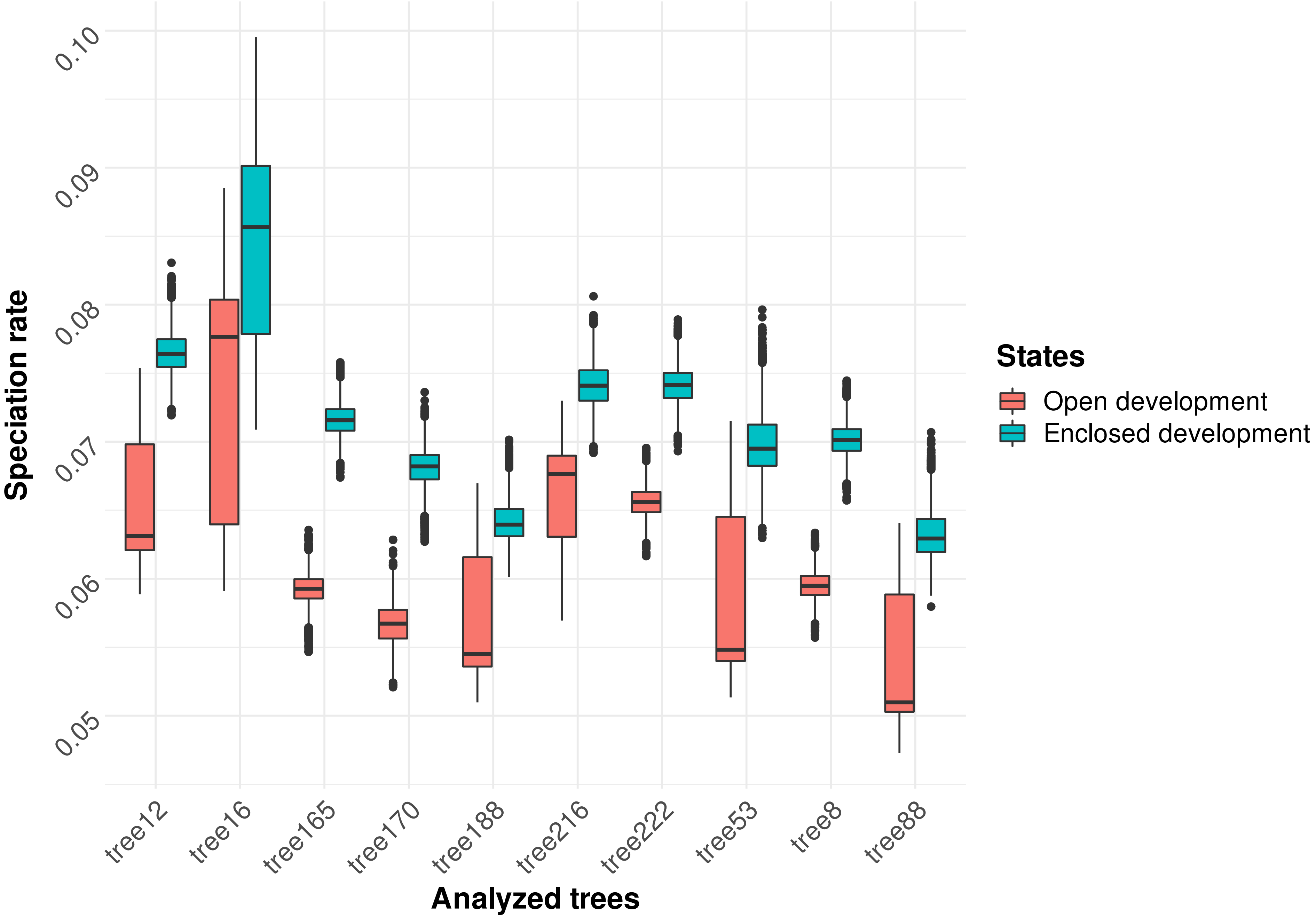
