## Supplementary material for "Developmental innovations promote species diversification in mushroom-forming fungi": figure S3

### Agaricoid clade sensu Matheny et al. 2016

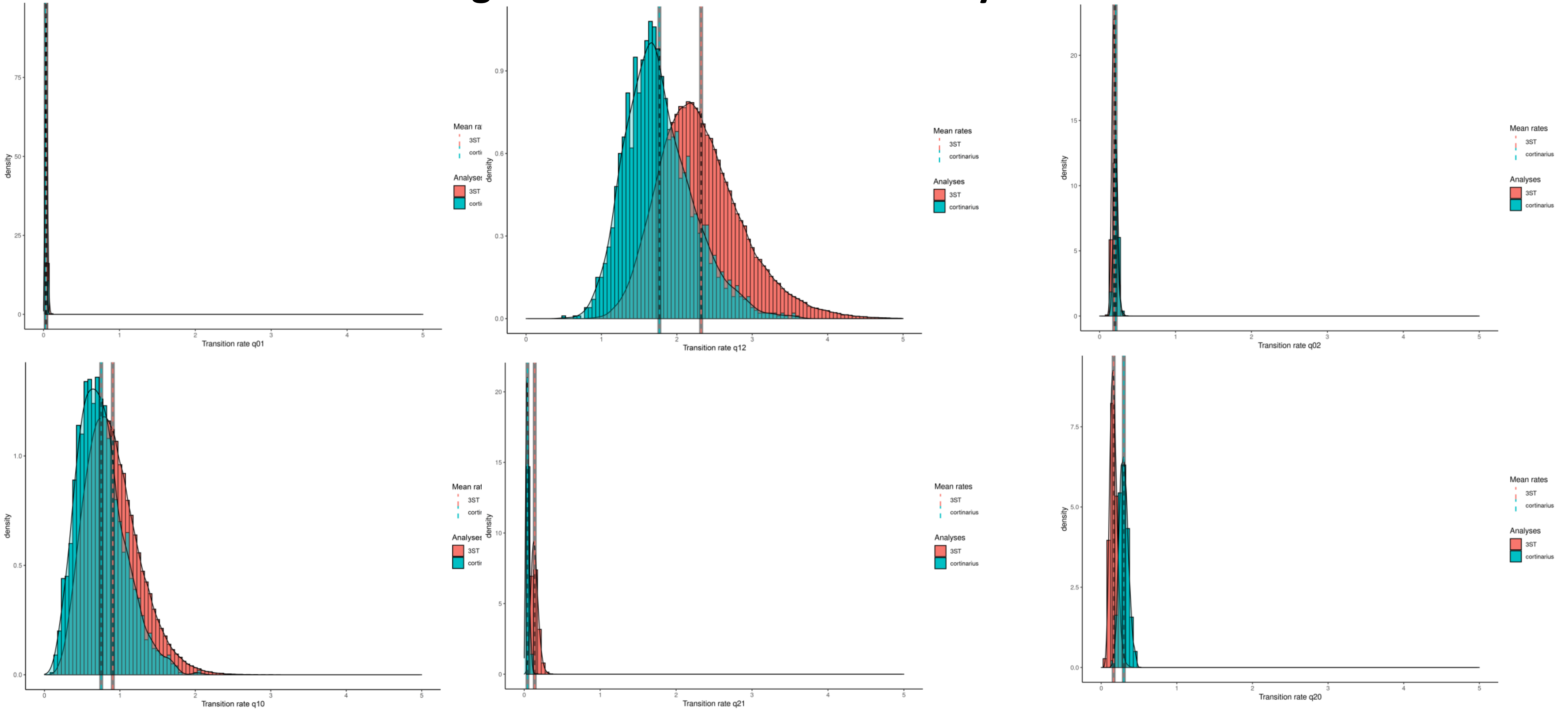


| Coding regime | q01 | | q10 | | q12 | | q21 | | q02 | | q20 | |
|---|---|---|---|---|---|---|---|---|---|---|---|---|
| | Mean | SD | Mean | SD | Mean | SD | Mean | SD | Mean | SD | Mean | SD |
| 3ST | 0.04 | 0.01 | 0.91 | 0.35 | 2.32 | 0.57 | 0.14 | 0.05 | 0.19 | 0.03 | 0.16 | 0.05 |
| 3ST Agaricoid clade constrained | 0.03 | 0.01 | 0.75 | 0.31 | 1.77 | 0.45 | 0.04 | 0.02 | 0.21 | 0.03 | 0.30 | 0.06 |

### Marasmioid clade sensu Matheny et al. 2016

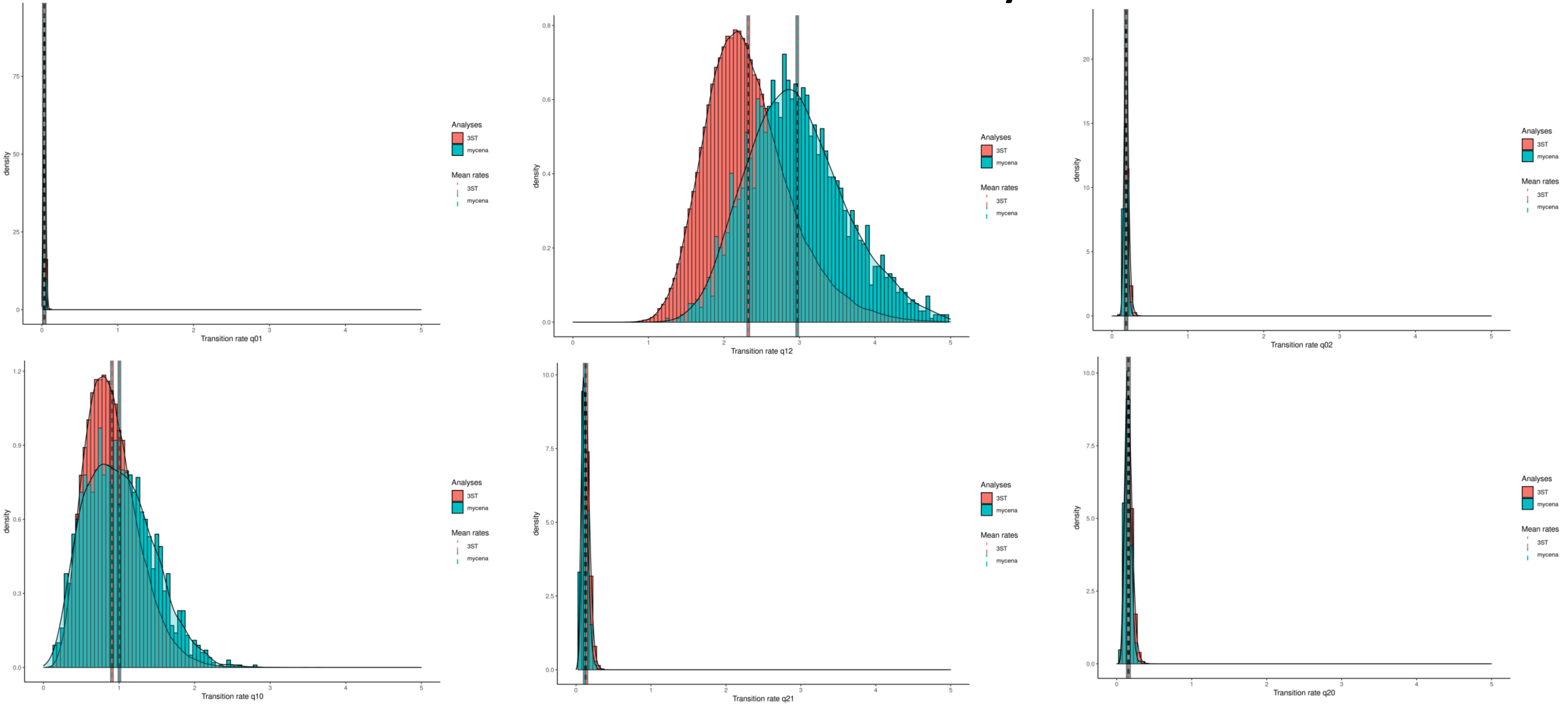


| Coding regime | q01 | | q10 | | q12 | | q21 | | q02 | | q20 | |
|---|---|---|---|---|---|---|---|---|---|---|---|---|
| | Mean | SD | Mean | SD | Mean | SD | Mean | SD | Mean | SD | Mean | SD |
| 3ST | 0.04 | 0.01 | 0.91 | 0.35 | 2.32 | 0.57 | 0.14 | 0.05 | 0.19 | 0.03 | 0.16 | 0.05 |
| 3ST Marasmioid clade constrained | 0.03 | 0.01 | 1.00 | 0.44 | 2.97 | 0.66 | 0.11 | 0.04 | 0.18 | 0.03 | 0.15 | 0.04 |

### Tricholomatoid clade sensu Matheny et al. 2016

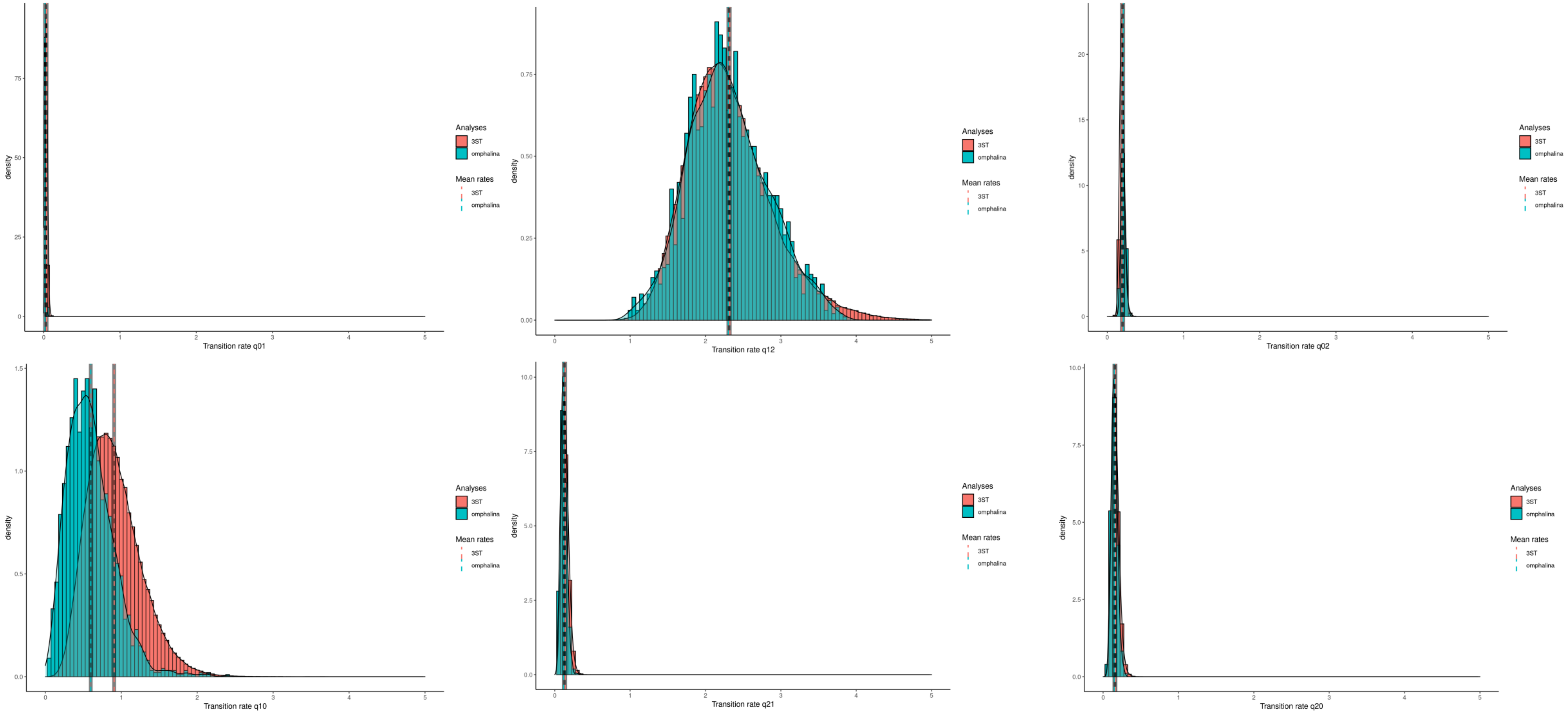


| Coding regime | q01 | | q10 | | q12 | | q21 | | q02 | | q20 | |
|---|---|---|---|---|---|---|---|---|---|---|---|---|
| | Mean | SD | Mean | SD | Mean | SD | Mean | SD | Mean | SD | Mean | SD |
| 3ST | 0.04 | 0.01 | 0.91 | 0.35 | 2.32 | 0.57 | 0.14 | 0.05 | 0.19 | 0.03 | 0.16 | 0.05 |
| 3ST Tricholomatiod clade constrained | 0.02 | 0.01 | 0.60 | 0.31 | 2.30 | 0.53 | 0.12 | 0.04 | 0.21 | 0.03 | 0.15 | 0.04 |

### Psathyrellaceae

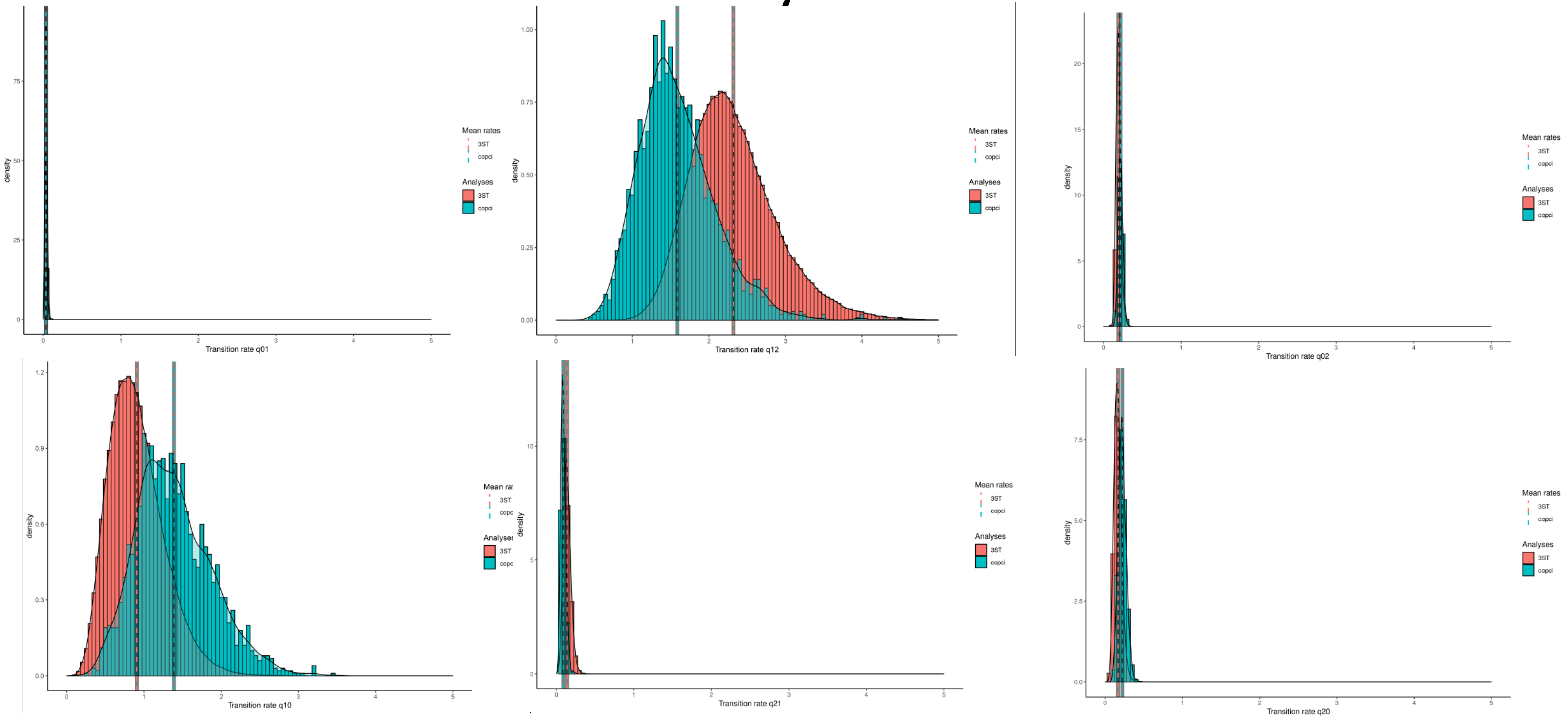


| Coding regime | q01 | | q10 | | q12 | | q21 | | q02 | | q20 | |
|---|---|---|---|---|---|---|---|---|---|---|---|---|
| | Mean | SD | Mean | SD | Mean | SD | Mean | SD | Mean | SD | Mean | SD |
| 3ST | 0.04 | 0.01 | 0.91 | 0.35 | 2.32 | 0.57 | 0.14 | 0.05 | 0.19 | 0.03 | 0.16 | 0.05 |
| 3ST Psathyrellaceae constrained | 0.03 | 0.01 | 1.39 | 0.49 | 1.59 | 0.50 | 0.14 | 0.04 | 0.22 | 0.03 | 0.22 | 0.05 |

### Boletales

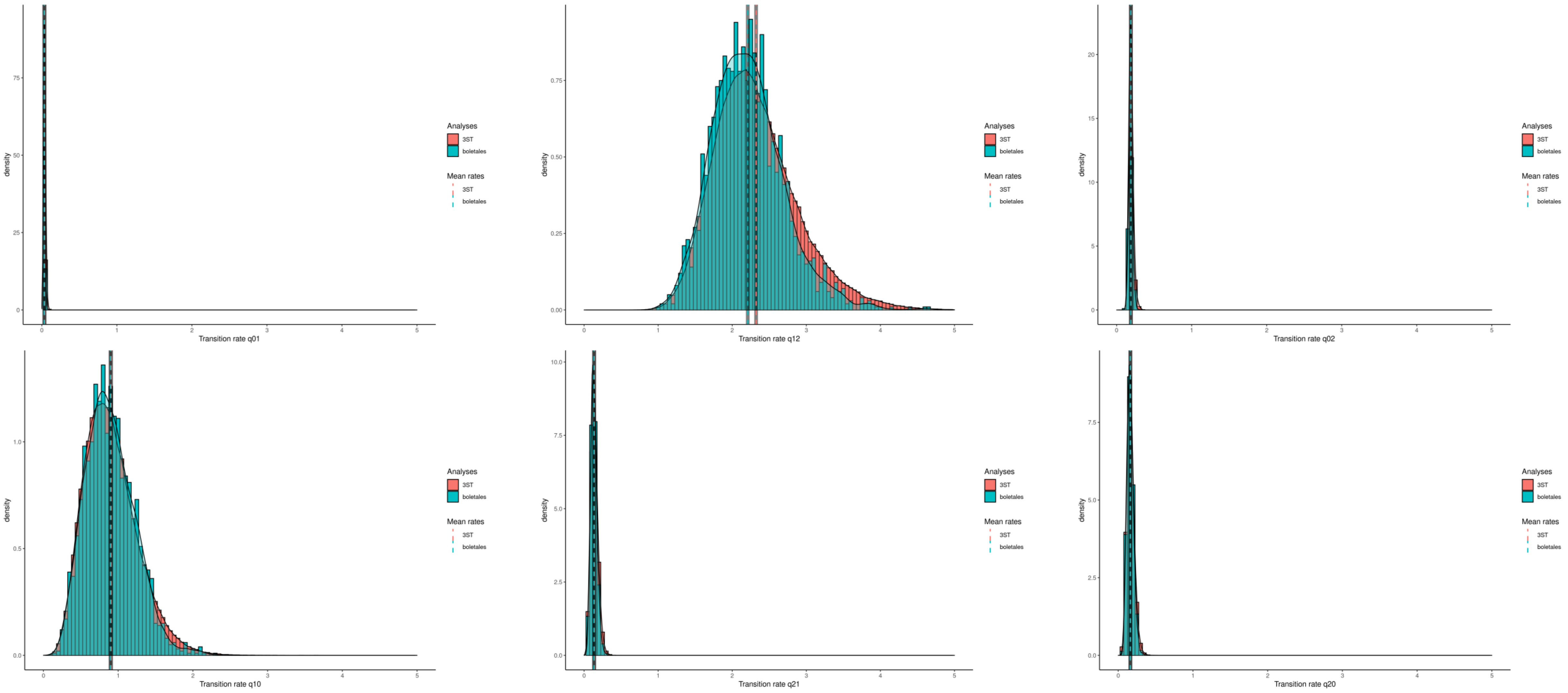


| Coding regime | q01 | | q10 | | q12 | | q21 | | q02 | | q20 | |
|---|---|---|---|---|---|---|---|---|---|---|---|---|
| | Mean | SD | Mean | SD | Mean | SD | Mean | SD | Mean | SD | Mean | SD |
| 3ST | 0.04 | 0.01 | 0.91 | 0.35 | 2.32 | 0.57 | 0.14 | 0.05 | 0.19 | 0.03 | 0.16 | 0.05 |
| 3ST Boletales constrained | 0.03 | 0.01 | 0.90 | 0.33 | 2.21 | 0.49 | 0.13 | 0.04 | 0.19 | 0.03 | 0.16 | 0.04 |

### Russulales

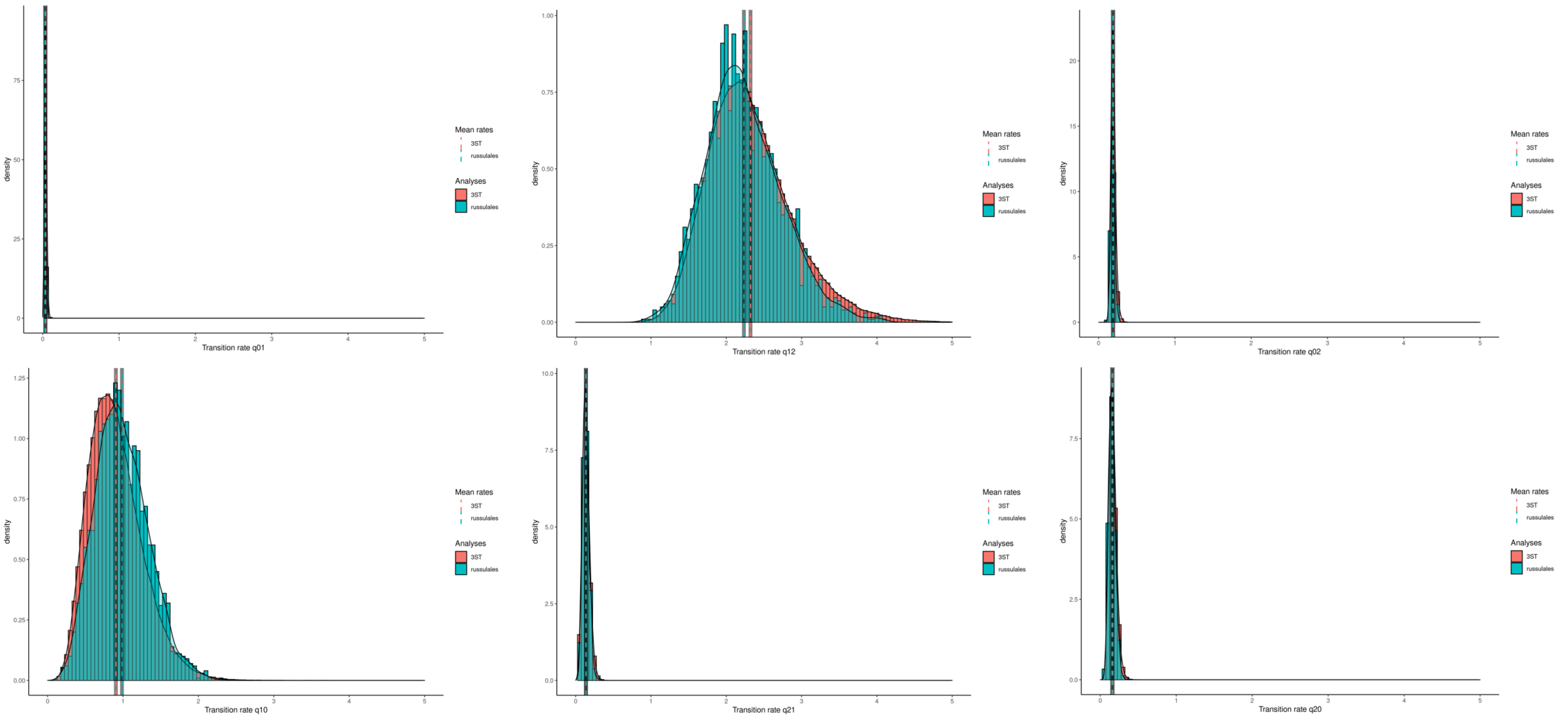


| Coding regime | q01 | | q10 | | q12 | | q21 | | q02 | | q20 | |
|---|---|---|---|---|---|---|---|---|---|---|---|---|
| | Mean | SD | Mean | SD | Mean | SD | Mean | SD | Mean | SD | Mean | SD |
| 3ST | 0.04 | 0.01 | 0.91 | 0.35 | 2.32 | 0.57 | 0.14 | 0.05 | 0.19 | 0.03 | 0.16 | 0.05 |
| 3ST Russulales constrained | 0.03 | 0.01 | 0.99 | 0.35 | 2.23 | 0.50 | 0.14 | 0.04 | 0.19 | 0.03 | 0.16 | 0.04 |
