## Supplementary material for "Developmental innovations promote species diversification in mushroom-forming fungi": text S1

**Supplement text**

**Scrutiny of inferred transition rates of enclosed development**

We inferred high transition rates from semi-enclosed development towards open and enclosed development (see the main text). Here, we performed series of analyses to check whether this pattern still holds under alternative character coding regimes.

We analyzed four alternative character coding regimes along three lines of thinking. First, we tested whether highly derived fruiting body morphologies disproportionately drove the inferred patterns by recoding them as alternative character states (see methods). This resulted in four different coding regimes (4ST1, 4ST2, 4ST3, and 5ST). Second, given the difficulty of recognizing the semi-enclosed state, we addressed whether lumping it together with other states impacts our inferences. Here, we produced three character state coding regimes by merging semi-enclosed and enclosed development states (2ST1), or open and semi-enclosed development (2ST2) or randomly recoding semi-enclosed states to either open and enclosed development states (2ST3). Third, we wanted to uncover if a particular part of or phylogeny drives the observed pattern. To do this, we examined the effects of character states of basal nodes and major clades.

We analyzed three four-character-state datasets (4ST1, 4ST2, 4ST3) to explore the impact of recoding gasteroid and secotioid species as a separate state and that of the coding of several hard-to-assign species (see Methods). With this character state coding, we tested whether the sporadic distribution of gasteroid could affect the transition rate patterns. We also treated ambiguous character states differently among the three four-states coding regimes to uncover the possible effects of ambiguous character states. A five-state coding (5ST) was also created by assigning a fifth character state to cyphelloid species in the 4ST1 coding regime. This addressed whether the mosaic distribution of cyphelloid species, which evolved via secondary simplification (Bodensteiner et al. 2004, Varga et al. 2019) could cause frequent backward transitions from semi-enclosed to open development. We found that the transition rate pattern observed above (high q10 and q12 and low q21 and q01) was robust and consistent through all alternative coding regimes (Supplementary Table 2., Supplementary Table 3).

Next, we hypothesized that the early evolution of semi-enclosed development could cause frequent transition towards open development. Therefore, we rerun MCMC analyses with 3ST coding regime while certain basal nodes were fixed (see materials and methods). We found that constraining 12 ancestral nodes to be open developmental states did not affect the transition rate pattern described above. (Supplementary Fig. 2.).

Finally, we tested if the coding of any major clade has a relevant effect on the transition rate values. We modified state 1 and state 01 to state 0 and state 12 to state 2 in each of the Marasmioid, Tricholomatoid, Agaricoid clades (sensu Matheny et al. 2006), as well as the Psathyrellaceae, Boletales and Russulales, producing six different coding regimes (Supplementary Table 1.). These changes did not appreciably affect the inferred transition rates (up to a mean transition rate difference of 0 – 0.11 relative to the 3ST analyses), except the Marasmioid and Agaricoid clades caused somewhat larger changes in the transition rates (up to 0.65). Nevertheless, the relative ratios of transition rates (high q_10_ and q_12_ and low q_01_ and q_21_) were consistent in all cases (Supplementary Fig. 3.).

Overall, we found that the evolution of enclosed development could go through the semi-enclosed development or could directly evolve from open development, and we demonstrated that the transition rate pattern we described above is robust and valid given our data.

**Analyzing the speciation and extinction rate of enclosed development 2ST1, 2ST2 and 2ST3 regimes**

We also analyzed two-state datasets where the semi-enclosed development was lumped together with other states. These analyses were performed on trait dependent diversification model (BiSSE) to assess the changes in sate specific speciation and extinction rates. When semi-enclosed development and enclosed development states were lumped together (2ST1), we found that the reverse transition rate (q_10_) was 1.1 times higher than the forward transition rate (q_01_), which proved to be insignificant differences according to likelihood ratio tests (LRT, p < 0.05, Supplementary Table 5.). The differences between the two rates were slightly increased when the semi-enclosed and the open development was lumped together (2ST2, q_10_/q_01_ = 1.5), which was a significant change in four trees out of the ten (LRT, p < 0.05, Supplementary Table 5.). The transition rate asymmetry was the highest (q_10_/q_01_ = 3.5) when two states coding was created by randomly assigning open or enclosed development to species which was initially coded as semi-enclosed development (2ST3). Speciation rates were relatively robust to the different coding regimes. We found that the speciation rate of state 1 was 1.16 - 1.24 times higher than that of state 0 depend on coding regimes. These differences were significantly different according to the LRT across all analyses and trees (p < 0.05).

**References**

Bodensteiner, P., Binder, M., Moncalvo, J. M., Agerer, R., & S. Hibbett, D. (2004). Phylogenetic relationships of cyphelloid homobasidiomycetes. *Molecular Phylogenetics and Evolution*, *33*(2), 501–515. https://doi.org/10.1016/j.ympev.2004.06.007

Varga, T., Krizsán, K., Földi, C., Dima, B., Sánchez-García, M., Sánchez-Ramírez, S., Szöllősi, G. J. G. J., Szarkándi, J. G. J. G., Papp, V., Albert, L., Andreopoulos, W., Angelini, C., Antonín, V., Barry, K. W. K. W., Bougher, N. L. N. L., Buchanan, P., Buyck, B., Bense, V., Catcheside, P., … Nagy, L. G. L. G. (2019). Megaphylogeny resolves global patterns of mushroom evolution. *Nature Ecology & Evolution*, *3*(4), 668–678. https://doi.org/10.1038/s41559-019-0834-1
